# Neurexin and Frizzled intercept axonal-transport at microtubule minus-ends to control synapse formation

**DOI:** 10.1101/2021.03.22.436477

**Authors:** Santiago Balseiro-Gómez, Junhyun Park, Yang Yue, Chen Ding, Lin Shao, Selim Ҫetinkaya, Caroline Kuzoian, Marc Hammarlund, Kristen J Verhey, Shaul Yogev

## Abstract

Precise synaptic connectivity defines neuronal circuits. Synapse formation is locally determined by transmembrane proteins, yet synaptic material is synthesized remotely and undergoes processive transport in axons. How local synaptogenic signals intercept synaptic cargo in transport to promote its delivery and synapse formation is unknown. We found that control of synaptic cargo delivery at microtubule (MT) minus-ends mediates pro- and anti-synaptogenic activities of presynaptic Neurexin and Frizzled in *C. elegans*, and identified the atypical kinesin VAB-8/KIF26 as a key molecule in this process. VAB-8/KIF26 levels at synaptic MT minus-ends are controlled by Frizzled and Neurexin, its loss mimics neurexin mutants or Frizzled hyperactivation, and its overexpression can rescue synapse-loss in these backgrounds. VAB-8/KIF26 is required for the synaptic localization of other minus-end proteins and promotes pausing of retrograde transport to allow delivery to synapses. Consistently, reducing retrograde transport rescues synapse-loss in *vab-8* and neurexin mutants. These results uncover an important mechanistic link between synaptogenic signaling and axonal transport.

## Introduction

The proper function of neuronal circuits hinges on establishment and maintenance of precisely patterned synaptic connections. Synapse position and numbers are instructed by local pro- and anti-synaptogenic signals from adhesion molecules such as Neurexin/Neuroligin and diffusible cues such as Wnts and their Frizzled (Fz) receptors (Sanes and Zipursky, 2020; Südhof, 2018; Yogev and Shen, 2014). However, most synaptic proteins are synthesized in the cell-body and rely on long-range transport to arrive at synapses (Guedes-Dias and Holzbaur, 2019; Maeder et al., 2014a; Rizalar et al., 2021). A key question is therefore how the activity of synaptogenic signals locally controls the delivery of cargo undergoing transport.

Presynaptic Neurexins are key synapse-organizing molecules, with variable context-dependent effects on synapse formation and maturation. Neurexin’s extra-cellular domain binds postsynaptic ligands to specify synapse properties (Südhof, 2017). Despite the identification of Neurexin intracellular domain binding proteins, signaling events downstream of presynaptic neurexin are poorly understood (Biederer and Südhof, 2000; Brouwer et al., 2019; Hata et al., 1996; Owald et al., 2012). Wnts and their Fz receptors are universal signaling molecules with diverse developmental and homeostatic roles. Unlike Neurexins, which promote synapse formation or specialization, Fz can either promote or inhibit synaptogenesis (He Chun-Wei et al.; Koles and Budnik, 2012; Sanes and Zipursky, 2020). Both Fz and Neurexin can signal to the synaptic cytoskeleton, although how this regulates synapse formation is poorly understood (Biederer and Sudhof, 2001; Lüchtenborg et al., 2014; Miech et al., 2008; Muhammad et al., 2015; Sugie et al., 2015).

Synapses rely on microtubule (MT)-based transport of cargo (Kreutzberg, 1969). Kinesin-3/KIF1A/UNC-104 is the key motor for delivering synaptic cargo (Hall and Hedgecock, 1991; Okada et al., 1995; Pack-Chung et al., 2007). However, live-imaging clearly shows that cargo is also delivered to *en passant* synapses during retrograde transport, suggesting a role for the retrograde motor dynein in synapse formation (Bharat et al., 2017; Wong et al., 2012). While several mechanisms that locally promote capture of dense core vesicles (DCVs) transported by Kif1A/kinesin-3 at synaptic sites were identified (Bharat et al., 2017; Stucchi et al., 2018), how cargo capture during retrograde transport is regulated is largely unknown.

Dynamic MT plus-ends are abundant at synapses and were recently shown to locally promote SVP delivery during anterograde transport by kinesin-3 (Guedes-Dias et al., 2018; Qu et al., 2019). Retrograde transport preferentially pauses at MT minus-ends (Soundararajan and Bullock, 2014; Tan et al., 2018; Yogev et al., 2016), raising the possibility that dynein-driven transport is regulated at these sites.

Here we found that control of synapse number and positioning by *C. elegans* Neurexin (NRX-1) and the Fz receptor MIG-1 involve cargo delivery at MT minus-ends. We identified the atypical kinesin VAB-8/KIF26 as a MT minus-end resident protein that functions in this process. NRX-1 and Fz signaling play antagonistic roles in controlling VAB-8/KIF26 levels on a subset of synaptic MTs, which correlate with their pro – and anti – synaptogenic functions, respectively. Local loss of VAB-8 from synaptic MT minus-ends results in impaired distribution of other MT minus-end proteins PTRN-1/CAMSAP and NOCA-1/Ninein, and in excessively processive retrograde transport, which leads to synapse loss. Consistently, reducing dynein activity restores the pattern of synapses in *vab-8*/KIF26 and *nrx-1* mutants. These results uncover cargo delivery at MT minus-ends as a mechanism for sculping synaptic connectivity by Neurexin and Frizzled signaling.

## Results

### *shy20* and *shy8* are required for synapse formation

The *C. elegans* cholinergic motor neuron DA9 resides in the preanal ganglion, from which it extends a dendrite in the ventral cord and an axon that traverses to the dorsal cord and grows anteriorly. Following the commissure is a short asynaptic domain, followed by ∼30 *en passant* presynaptic sites at stereotypic positions (Figure 1A) (White et al., 1986). Expression of the active zone protein CLARINET-1 (CLA-1) and the SVP marker RAB-3 allow the visualization of DA9 synapses at single-cell resolution (Chen et al., 2018; Kurshan et al., 2018; Xuan et al., 2017) (Figure 1B).

**Figure 1:**
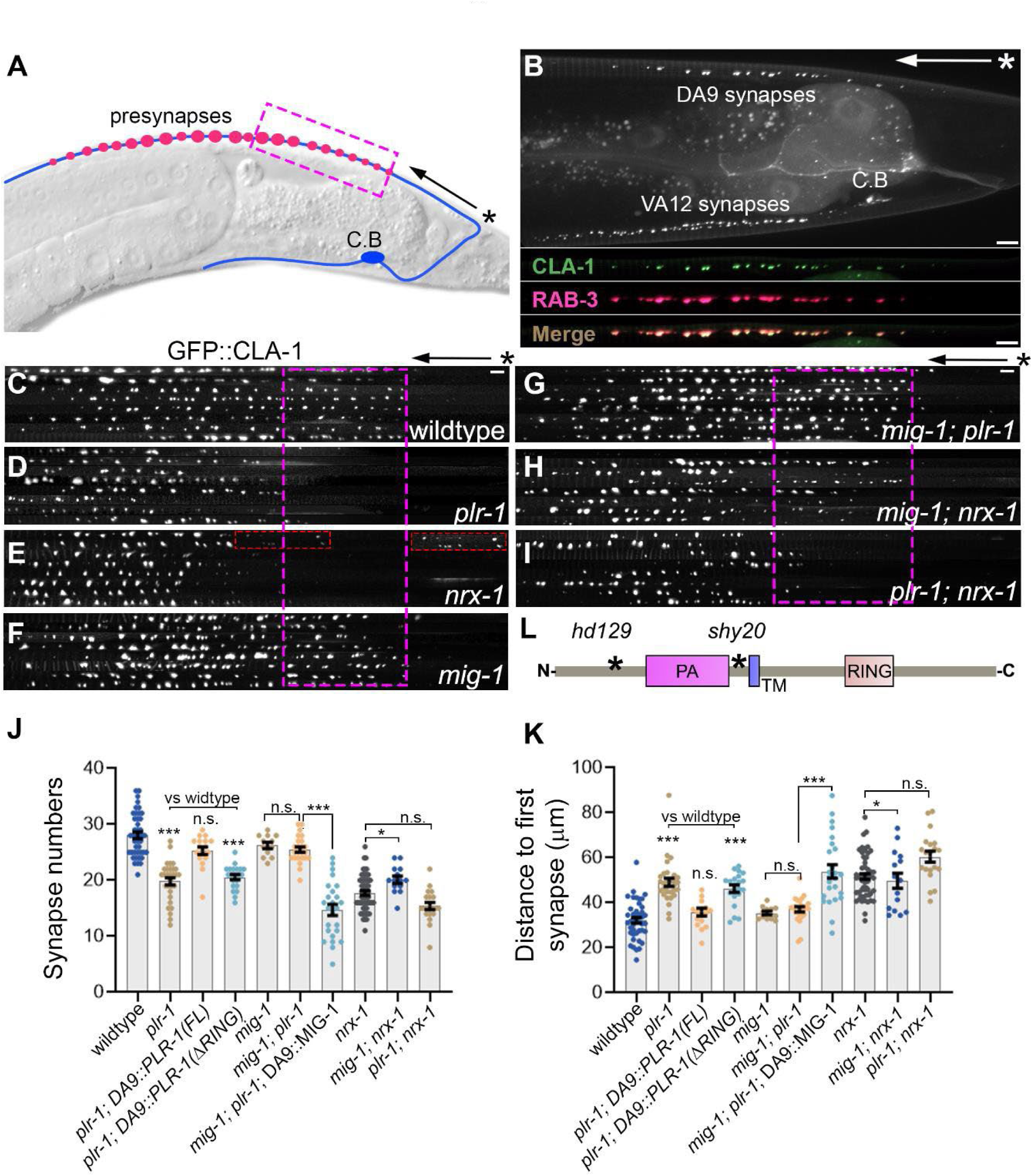
Fz/MIG-1 signaling acts antagonistically to Neurexin in synapse formation. (A) Schematic of the DA9 motor neuron. ∼30 en passant presynapses form at stereotypic position along the axon. * and arrow indicate the turn of the commissure and the asynaptic domain in all figures. Magenta brackets indicate the region affected in *vab-8* and *plr-1* mutants. (B) wildtype adult expressing the active zone marker GFP::CLA-1 and the synaptic vesicle marker tdTomato::RAB-3 under the P*mig-1*3 promoter. VA12 synapses on the ventral side are also labelled by the transgene. Bottom panels show a straightened section of the axon beginning at the turn of the commissure (*). C.B, DA9 cell body. Scale bar 5 ⍰m. (C-I) Confocal images of wildtype (C) and indicated mutant genotypes expressing the synaptic marker GFP::CLA-1. Eight axons per genotype were straightened and aligned from the turn of the commissure (*). Magenta square highlights the missing synapses in *plr-1* and *nrx-1* mutants. Inset in (E) shows fainter proximal CLA-1 puncta in *nrx-1* mutants. These were counted as synapses if their intensity was >5% of the brightest CLA-1 peak. Scale bar 5 μm. (J, K) Quantifications of synapse numbers (j) and distance to first synapse (K) in indicated genotypes. n=15-54 ***p<0.001; **p<0.01; *p<0.05 (Mann-Whitney U test). (L) Domain structure of PLR-1 showing *shy20* and hd129 allele. PA, protease-associated domain. TM, transmembrane domain. RING, RING finger.

In a genetic screen for MT regulators, we found that two mutants, *shy20* and *shy8*, displayed a specific loss of proximal synapses (Figures 1,2, S1). This phenotype was highly reminiscent of *nrx-1 (wy778)* mutants (Kurshan et al., 2018) (For description of alleles, refer to Table S1. All alleles are strong loss of function or nulls, unless otherwise noted). The sole *C. elegans* Neurexin, *nrx-1*, is required presynaptically to cluster RAB-3 vesicles and prevent loss of proximal synaptic boutons (Kurshan et al., 2018) (Figure 1E). RAB-3 vesicle clustering around active zones was normal in *shy20* and *shy8* mutants (Figure S1C, D), suggesting that they may be specifically related to Neurexin’s function in defining synapse number and position rather than vesicle clustering. Synapse spacing, CLA-1 or RAB-3 puncta size, and the length of the synaptic domain were largely normal in *shy20* and *shy8* mutants (Figure S1F-I). We observed a clear negative correlation between synapse numbers and the distance from the commissure to the proximal-most bouton in all three mutants (Figure S1E). We therefore used these two parameters in subsequent quantifications of interactions between *shy20*, *shy8*, and *nrx-1*.

**Figure 2:**
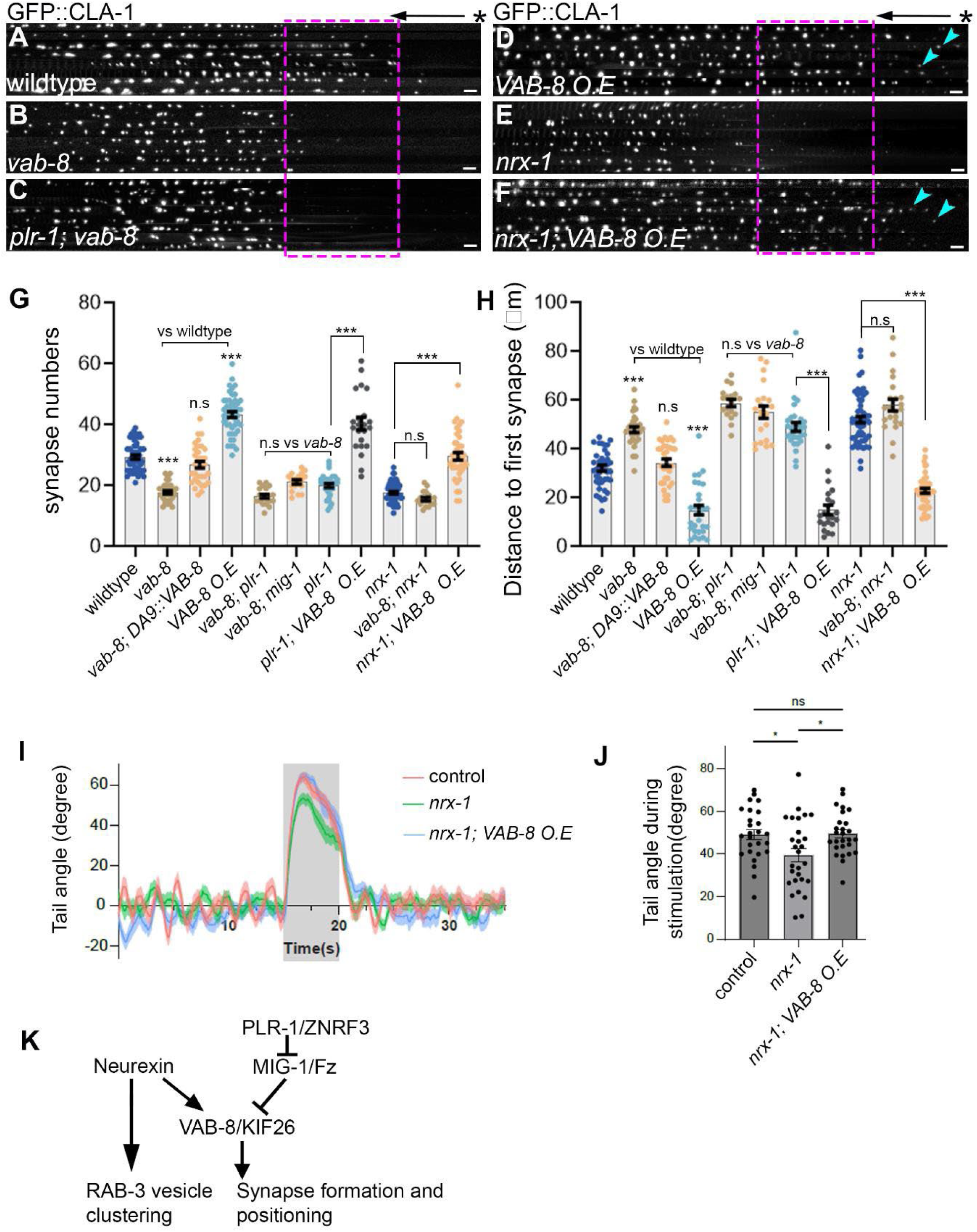
The atypical kinesin VAB-8/KIF26 controls synapse formation downstream of NRX-1/Neurexin and MIG-1/Fz. (A-E) Alignments of 8 axons expressing GFP::CLA-1 from confocal images of the following genotypes: wildtype (A), *vab-8* (B), *plr-1*; *vab-8* (C), Punc-17::VAB-8 (D), *nrx-1* (E), *nrx-1*, Punc-17::VAB-8 (F). (*) marks turn of the commissure and arrow marks the asynaptic region. Scale bar = 5 μm. Arrowheads in (D) and (F) indicate ectopic CLA-1 puncta induced by VAB-8 overexpression. (G-H) Quantification of synapse numbers (G) and the distance from the turn of the commissure to the first synapse (H). n = 15-54, ***p<0.001; **p<0.01; *p<0.05 (Mann-Whitney U test). (I) DA9 neurons expressing the Channel Rhodopsin Chrimson were activated for 5 seconds (grey shading) and the degree of tail bending induced by stimulation was measured. (J) Quantification of tail angle induced by optogenetically stimulating DA9 in indicated genotypes. n=22 per genotype *p<0.05 (Mann-Whitney U test). (K) Summary of genetic interactions among presynaptic *nrx-1*, *vab-8*, *plr-1* and *mig-1*. See text for details.

DA9 forms dyadic synapses onto ventral muscles and inhibitory GABA neurons (White et al., 1986). To test whether postsynaptic structures are lost in *shy8* and *shy20* mutants, we examined the cholinergic receptor subunit ACR-12::GFP (Petrash et al., 2013), expressed under a GABA promoter. ACR-12::GFP overlapped with a presynaptic marker, RFP::RAB-3, expressed in DA9 (Figure S2A-C). In *shy8* and *shy20* mutants we observed loss of postsynaptic ACR-12::GFP puncta at the same region where presynaptic markers are lost in these mutants (Figure S2C-H). We conclude that *shy20* and *shy8* are required for synapse formation in DA9, particularly at the proximal region.

### The Fz receptor MIG-1 acts antagonistically to Neurexin in synapse formation

We mapped *shy20* using whole-genome sequencing and rescue experiments to a premature stop codon (R188>Opal) in the transmembrane E3 ubiquitin ligase PLR-1, homolog of mammalian RNF43 and ZNRF3 (Figure 1L). A second allele, *plr-1(hd129)* showed a similar phenotype to *shy20* (not shown). Using rescue experiments, we determined that PLR-1 functions cell-autonomously in DA9 and that its function requires the RING domain, consistent with its role as a ubiquitin ligase (Figure 1 J, K). Double mutants between a *nrx-1 (wy778)* and *plr-1 (shy20)* did not enhance the loss of synapses observed in single mutants, suggesting that *plr-1* and *nrx-1* function in a common genetic pathway (Figure 1I, J, K).

PLR-1 and its mammalian homologs ubiquitinate Frizzled (Fz) receptors to downregulate signaling (Hao et al., 2012; Koo et al., 2012; Moffat et al., 2014). In DA9, the Fz receptor LIN-17 mediates anti-synaptogenic functions of its ligand LIN-44, but also promotes synapse-formation independently of LIN-44 (Klassen and Shen, 2007; Kurshan et al., 2018). We therefore speculated that synapse loss in *plr-1* mutants was mediated by LIN-17. However, *lin-17*; *plr-1* double mutants were similar to *plr-1* mutants, indicating that LIN-17 does not function downstream of PLR-1 (Figure S3D). Consistently, we did not observe changes in the subcellular distribution of LIN-17::YFP in *plr-1* mutants (Figure S3G). We next examined the Fz receptor MIG-1, which does not have a known function in DA9. A MIG-1::GFP transgene expressed in DA9 was enriched in the dendrite but also localized to the axon. In the axon, MIG-1::GFP localized to the region where synapses are lost in *plr-1* mutants (Figure S3H). This localization is consistent with the presence of source cells secreting Wnt/EGL-20, MIG-1’s ligand, in this region, and with the ability of Wnt/EGL-20 to control MIG-1 localization (Mizumoto and Shen, 2013a; Pani and Goldstein, 2018). *mig-1* mutants suppressed synapse loss in *plr-1* mutants, strongly suggesting that PLR-1’s function is to downregulate MIG-1 signaling to prevent synapse loss (Figure 1G, J, K). This suppression was reversed by expression of a MIG-1 transgene in DA9, indicating that MIG-1 functions cell-autonomously (Figure 1 J, K).

We next examined if *mig-1* loss-of-function can also rescue *nrx-1* mutants. In *mig-1*; *nrx-1* double mutants, synapse loss was rescued compared to *nrx-1* single mutant, although only partially (Figure 1H, J, K). These results indicate that MIG-1 signaling acts either downstream or in parallel to Neurexin to antagonize synapse formation. Since NRX-1 also has MIG-1/Fz independent functions, such as RAB-3 vesicle clustering, we propose that it functions in parallel, rather than upstream, to MIG-1/Fz (Figure 2K).

### The atypical kinesin VAB-8/KIF26 is required for synapse formation downstream of NRX-1/Neurexin and MIG-1/Fz

The second mutant which displayed a loss of proximal synapses, *shy8*, mapped to VAB-8, an atypical kinesin homologous to mammalian KIF26. Kif26A functions in enteric and sensory neurons, where it regulates GDNF-Ret and FAK signaling, respectively. Kif26B regulates cell polarization and migration (Guillabert-Gourgues et al., 2016; Susman et al., 2017; Wang et al., 2018; Zhou et al., 2009) and mutations in human KIF26B are associated with progressive microcephaly and spinocerebellar ataxia (Nibbeling et al., 2017; Wojcik et al., 2018). In *C. elegans*, *vab-8* is required for cell migration, axon extension of posteriorly growing neurons and GAP junction localization (Meng et al., 2016; Wightman et al., 1996; Wolf et al., 1998). DA9 grows anteriorly, and we did not detect DA9 axon growth or navigation defects in *vab-8* mutants or when VAB-8 was overexpressed (not shown).

*shy8* is a 153 bp deletion and frameshift leading to a stop codon shortly after the motor domain (Figure S1J). A CRISPR-induced stop codon in the same region *(shy72)* was indistinguishable from *shy8* and was used interchangeably. Comparison to previously isolated *vab-8* null alleles *(ev411, gm84, e1017)* or a trans-heterozygote with a genomic deletion encompassing the *vab-8* region *(shy8/arDf1)* revealed similar phenotypic severity (not shown and Figure S5 for MT phenotypes) suggesting that *shy8* is a null allele. Expression of VAB-8 in DA9 under the *Pmig-13* promoter rescued the Shy8 phenotype, revealing that VAB-8 functions cell-autonomously (Figure S1K,L, Figure 2G, H).

To gain insight into how VAB-8 functions, we assayed the requirement for its predicted domains in rescue experiments. Full length VAB-8 robustly rescued synapse loss in *vab-8* mutants, but constructs lacking the C-terminal coiled-coils or the N-terminal motor domain did not, consistent with VAB-8 function as a motor. A mutation in the motor domain (R279A), which reduces MT binding of kinesin-1 (Woehlke et al., 1997), resulted in reduced MT binding by the VAB-8 motor domain (Figure 3H, J) and rendered the VAB-8 transgene ineffective in rescuing *vab-8 (shy8)* mutants (Figure S1K-O). These results indicate that MT-binding is required for VAB-8 function.

**Figure 3:**
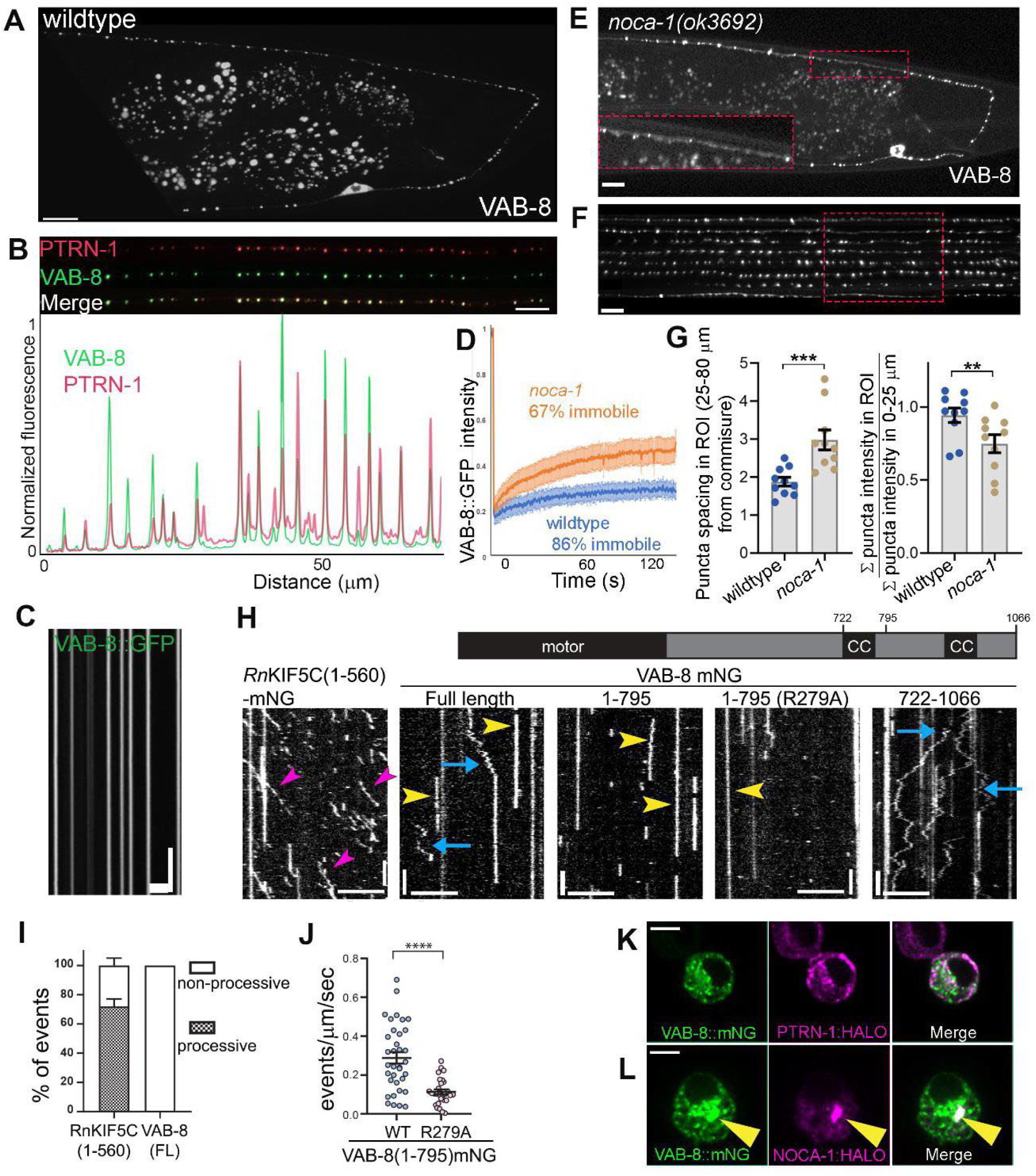
VAB-8 is a MT minus-end resident protein in neurons. (A) VAB-8::GFP in wildtype animals shows punctate staining in the axon and dendrite. Scale bars = 5 μm in all panels. (B) VAB-8::GFP co-localizes with the MT minus-end marker RFP::PTRN-1/CAMSAP. Lower panel shows normalized fluorescence intensity plots. (C) Kymograph from a movie of VAB-8::GFP showing no discernible movement in the axon. Scale = 5 μm and 10 sec. (D) FRAP traces of VAB-8::GFP in the DA9 axon in wildtype and *noca-1* mutants. n=7 per condition. (E) *noca-1* is required for the localization of VAB-8. VAB-8::GFP puncta frequency is mildly reduced in *noca-1*(ok3692) mutants derived from heterozygous animals (*noca-1* is required for viability in the embryo), with the strongest effect in proximal synapses. (F) Alignment of several axons from the same genotype as (E) shows that the region where VAB-8 loss occurs in *noca-1* mutants is most consistently the proximal synapses. (G) Quantification of the average spacing between VAB-8::GFP puncta in proximal synapses (25-80 μm from the turn of the commissure) and normalized puncta intensity in the same region (I) in wildtype and *noca-1* mutants. n = 10 per genotype ***p<0.001 (Mann-Whitney U test). (H) Kymographs from in vitro single-molecule motility assays of the indicated constructs on taxol-stabilized MTs. RnKIF5C (1-560) is a positive control for motile events (magenta arrowheads). Blue arrows indicated diffuse movement on the MT, and yellow arrowheads show immotile binding. Scale = 5 μm and 5 sec. (I) Quantification of processive and non-processive events (including diffusive, immotile and short MT binding events) of VAB-8 (n=102) compared to KIF5C control (n=187). (J) Quantification showing reduced MT binding caused by the R279A mutation in the motor domain. **** P<0.0001 two-tailed t-test, n=35 and 28 MTs (WT and R279A respectively. (K, L) S2 cells transfected with VAB-8::mNG and PTRN-1::HALO (K) or NOCA-1d::HALO (L). NOCA-1d can recruit VAB-8 to a MTOC-like structure. Scale bar = 5 μm.

To test whether VAB-8/KIF26 functions downstream of Neurexin and Fz signaling, we conducted double mutant analysis. *vab-8 (shy8)* mutants did not enhance synapse loss of *nrx-1 (wy778)* or *plr-1 (shy20)* mutants, suggesting that the three proteins function in a common genetic pathway (Figure 2B, C, G, H). Unlike the Plr-1 phenotype, which was suppressed by mutations in *mig-1*, *mig-1*; *vab-8* double mutants retained the synapse-loss phenotype of *vab-8* single mutants (Figure 2G, H). These results suggest that VAB-8 functions in parallel to MIG-1 or downstream to it, which is more consistent with results showing that MIG-1 reduces synaptic VAB-8 levels (see Figure 4).

**Figure 4:**
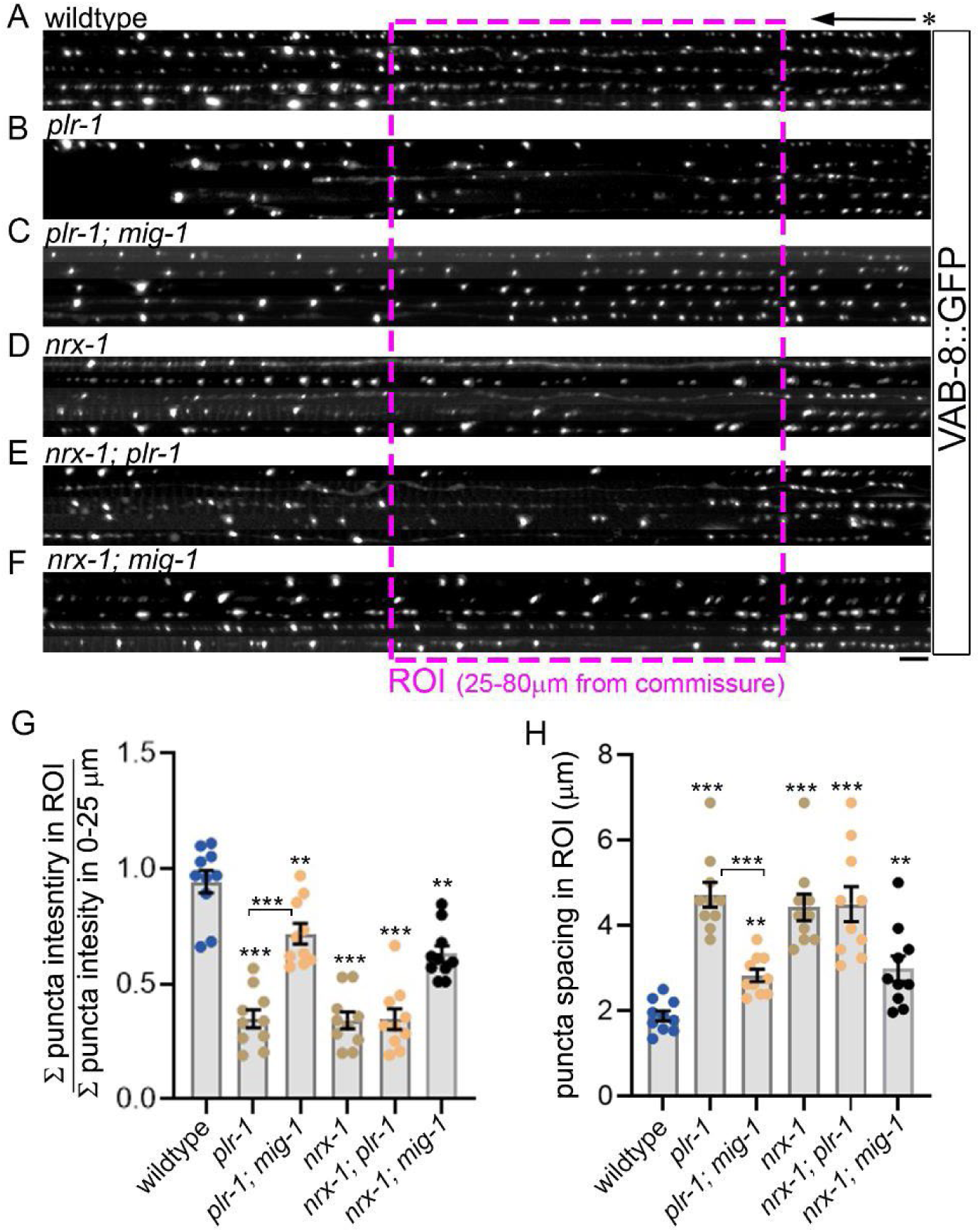
NRX-1/Neurexin and MIG-1/Fz control VAB-8 accumulation on synaptic MT minus-ends. (A-F). Confocal images of wildtype (A), *plr-1*(B), *plr-1*;*mig-1*(C), *nrx-1*(D), *nrx-1*;*mig-1*(E), *nrx-1*;*mig-1*(F) axons in DA9 expressing VAB-8::GFP. Five images from each genotype were straightened and aligned to the turn of the commissure (*). Scale bar = 5 μm. Magenta square highlights ROI around proximal synapses (25-80 μm from turn of commissure, similar to the region quantified in Figure 1) that was used in quantifications. (G) Quantification of VAB-8::GFP puncta intensity in proximal synapses (ROI) normalized by puncta intensity in the asynaptic domain (0-25 μm from commissure). (H) Spacing between VAB-8::GFP puncta in the region between 25-80 μm from the turn of the commissure. n=10 per genotype. ***p<0.001; **p<0.01; *p<0.05 (Mann-Whitney U test).

Next, we tested whether VAB-8 overexpression can compensate for loss of *plr-1* or *nrx-1*. Since overexpression using the *Pmig-13* promoter was lethal at high concentrations, we drove strong VAB-8 expression with a *Punc-17* cholinergic promoter. *Punc-17::VAB-8* fully rescued synapse loss in *plr-1* and *nrx-1* mutants (Figure 2F-H). Furthermore, VAB-8 overexpression in wildtype animals increased the numbers of presynaptic CLA-1 and RAB-3 clusters in the synaptic area and led to the formation of ectopic clusters in the asynaptic region (Figure 2D,G,H). These results indicate that VAB-8 promotes synapse formation in DA9 downstream of *plr-1* and *nrx-1*.

Overexpressing VAB-8 presynaptically (*Punc-17::VAB-8*) also rescued the loss of postsynaptic ACR-12::GFP puncta in *vab-8* and *plr-1* mutants (Figure S2G, H), confirming that loss of postsynapses in these mutants was due to presynaptic defects. Presynaptic Neurexin acts as a trans-synaptic organizer of postsynaptic ACR-12 in *C. elegans* (Philbrook et al., 2018). Consistently, we found that ACR-12::GFP was largely diffuse in *nrx-1* mutants, with very few dim puncta, even in opposition to normal presynaptic RAB-3 clusters. Although presynaptic VAB-8 overexpression modestly elevated ACR-12::GFP puncta number in *nrx-1* mutants, the overall dim and diffuse signal of ACR-12::GFP was not rescued. These results agree with VAB-8 acting downstream of Neurexin’s presynaptic, but not trans-synaptic function. We next used a behavioral assay to ask if the observed molecular rescue of synaptic markers by VAB-8 overexpression corresponded with improved overall synapse function. We optogenetically stimulated DA9 and measured the angle of the resulting dorsal bending of the worm’s tail (Ding and Hammarlund, 2018). *nrx-1* mutants showed a mild but consistent reduction in dorsal bending of the tail (Figure 2I, J).

The locomotion defects in *vab-8* mutants (flaccid posture, tail dragging and reduced antagonizing ventral bending, see Video S1) prevent a direct comparison with wildtype or *nrx-1* mutants in this assay. However, VAB-8 overexpression strongly rescued the phenotype of *nrx-1* mutants (Video S2, Figure 2I, J). Together with the rescue of synaptic markers, our data indicate that VAB-8 promotes synapse formation in DA9 downstream of Neurexin (Figure 2K).

### VAB-8 is a MT-minus-end resident protein in neurons

To understand how VAB-8 functions, we next examined its subcellular localization. To visualize VAB-8 at native levels in specific neurons, we inserted a 7xspGFP11 into the *vab-8* locus (He et al., 2019). When visualized with pan-neuronal spGFP1-10, VAB-8 appeared as distinct puncta with regular spacing in (Figure S4B). *vab-8* mRNA is detectable in DA9 (Figure S4A), but at levels which were not sufficient for visualizing the endogenous protein with a 7XspGFP11 tag. A functional VAB-8::GFP transgene expressed in DA9 (see rescue in Figure S4C-F) also showed punctate localization throughout the axon (Figure 3A), consistent with the distribution we observed for endogenous VAB-8 in other neurons. We noticed that the frequency of VAB-8::GFP puncta was similar to that of MT minus-ends in DA9 (Yogev et al., 2016). To test whether the VAB-8::GFP puncta we observed in the axon correspond to MT-minus ends, we co-expressed VAB-8::GFP with RFP::PTRN-1, which has been extensively validated as a minus-end marker (Feng et al., 2019; Goodwin and Vale, 2010; Jiang et al., 2014; Marcette et al., 2014; Meng et al., 2008; Wang et al., 2015). VAB-8::GFP puncta strongly colocalized with RFP::PTRN-1 puncta (Figure 3B), indicating that VAB-8 resides at minus-ends.

Most kinesins move processively to MT plus ends, so we asked how VAB-8 can localize to MT minus-ends. VAB-8 and KIF26 lack conserved residues for ATP hydrolysis in their motor domains. Consistently, KIF26 shows rigor-like co-sedimentation with MTs, suggesting that it is immotile (Terabayashi et al., 2012; Wolf et al., 1998; Zhou et al., 2009). We used single-molecule TIRF microscopy to image VAB-8::mNeonGreen (mNG) on Taxol-stabilized MTs in vitro and found that VAB-8::mNG was mostly immotile (Figure 3H,I). VAB-8::mNG can diffuse on and bind statically to the MT lattice but no directional movement was observed (Figure 3H). A truncated construct containing the VAB-8 motor domain (aa 1-795) was completely immotile, whereas a construct consisting of the C-terminal coiled coils (aa 722-1066) displayed mostly diffusive behavior (Figure 3H). Consistent with these results, live imaging of VAB-8::GFP in DA9 did not reveal directional motility (Figure 3C) and FRAP experiments showed an immobile fraction of ∼86% (Figure 3D).

Since VAB-8 was not enriched at the MT minus-end *in vitro* (Figure 3H), we tested whether known minus-end proteins recruit it to these sites *in vivo*. VAB-8::GFP maintained its punctate pattern in *ptrn-1* mutants, suggesting that *ptrn-1* is not required for VAB-8 localization to minus-ends (not shown). In the *C. elegans* germline and epidermis, a Ninein homolog, NOCA-1, localizes to minus-ends of non-centrosomal MTs and functions redundantly with PTRN-1. As *noca-1* mutants are sterile, we examined homozygous progeny of heterozygous mothers, which likely contain maternally contributed NOCA-1. In *noca-1* mutants VAB-8::GFP was reduced in the proximal synaptic region (Figure 3E, F, G). VAB-8::GFP was also more mobile in FRAP experiments in *noca-1* mutants compared to wildtype, suggesting that NOCA-1 helps to retain VAB-8 on minus-ends in the synaptic region. *ptrn-1; noca-1* double mutants did not further exacerbate the mislocalization of VAB-8::GFP (not shown). We next expressed VAB-8 with NOCA-1 or PTRN-1 in S2 cells to ask whether these proteins can influence VAB-8 localization. Expression of NOCA-1, but not PTRN-1, recruited VAB-8 to an MTOC-like structure (Figure 3K, L). Despite this effect, we could not detect a direct interaction between NOCA-1 and VAB-8 in vitro (not shown). Taken together, these results identify VAB-8 as a minus-end protein and suggest a role for NOCA-1/Ninein and unknown partner proteins in ensuring VAB-8 localization on synaptic MTs.

### NRX-1/Neurexin and MIG-1/Fz control VAB-8/KIF26 accumulation on synaptic MT minus-ends

Since loss of NOCA-1 reduced VAB-8::GFP puncta mostly in the synaptic domain, we asked whether local signals in that area control VAB-8 abundance. *plr-1* mutants, in which MIG-1/Fz signaling is increased, showed a strong loss of VAB-8::GFP in the proximal synaptic domain (Figure 4B, G). Loss of VAB-8 in *plr-1* mutants was more local and severe than in *noca-1* mutants (compare Figure 4G with 3H). Consistent with the synaptic phenotype, this effect could be significantly suppressed by loss of *mig-1* (Figure 4C, G), suggesting that the excessive MIG-1/Fz signaling in *plr-1* mutants disrupts VAB-8 localization. In *nrx-1* mutants, VAB-8 was also significantly reduced in the proximal synaptic region (Figure 4D, G). This reduction was not further enhanced in *plr-1*; *nrx-1* double mutants, suggesting that as in the case of synapse formation, both proteins function in the same genetic pathway. Loss of VAB-8::GFP in *nrx-1* mutants was partially dependent on MIG-1, since VAB-8::GFP was restored in *mig-1*; *nrx-1* mutants, although not completely (Figure 4F, G). The effects of *plr-1* and *nrx-1* on VAB-8 distribution on synaptic MT minus-ends are in agreement with the ability of overexpressed VAB-8 to suppress the phenotypes of *nrx-1* and *plr-1* mutants (Figure 2).

### VAB-8 is required for correct localization of PTRN-1 and NOCA-1 on synaptic MTs

We next examined how VAB-8 might function at MT minus ends. In wildtype animals, axonal MT minus-ends appear with regular periodicity and accumulate an overall uniform level of PTRN-1::YFP (Figure 5A, D). In *vab-8* and *plr-1* mutants, PTRN-1::YFP were sparser and dimmer at the proximal synaptic area, leading to a local “gap” in the signal (Figure 5B-F). This effect was observed in several alleles of *vab-8* and *plr-1* (Figure S5). Since in *plr-1* mutants VAB-8 is specifically reduced in the same region, these results, together with VAB-8 localization to minus-ends, argue that VAB-8 functions locally. Cell specific expression of VAB-8 and PLR-1 in DA9 rescued the phenotype, indicating that both proteins function cell-autonomously (Figure 5J). We also tested how loss of *vab-8* or *plr-1* affected NOCA-1/Ninein. NOCA-1::GFP showed a punctate distribution reminiscent of PTRN-1::YFP in wildtype animals (Figure 5G). In *vab-8* and *plr-1* mutants, NOCA-1::GFP was specifically reduced, but not completely eliminated, in the proximal synaptic domain (Figure 5G-I). These results indicate that VAB-8 is required for the normal distribution of other minus-end proteins on synaptic MTs.

**Figure 5.**
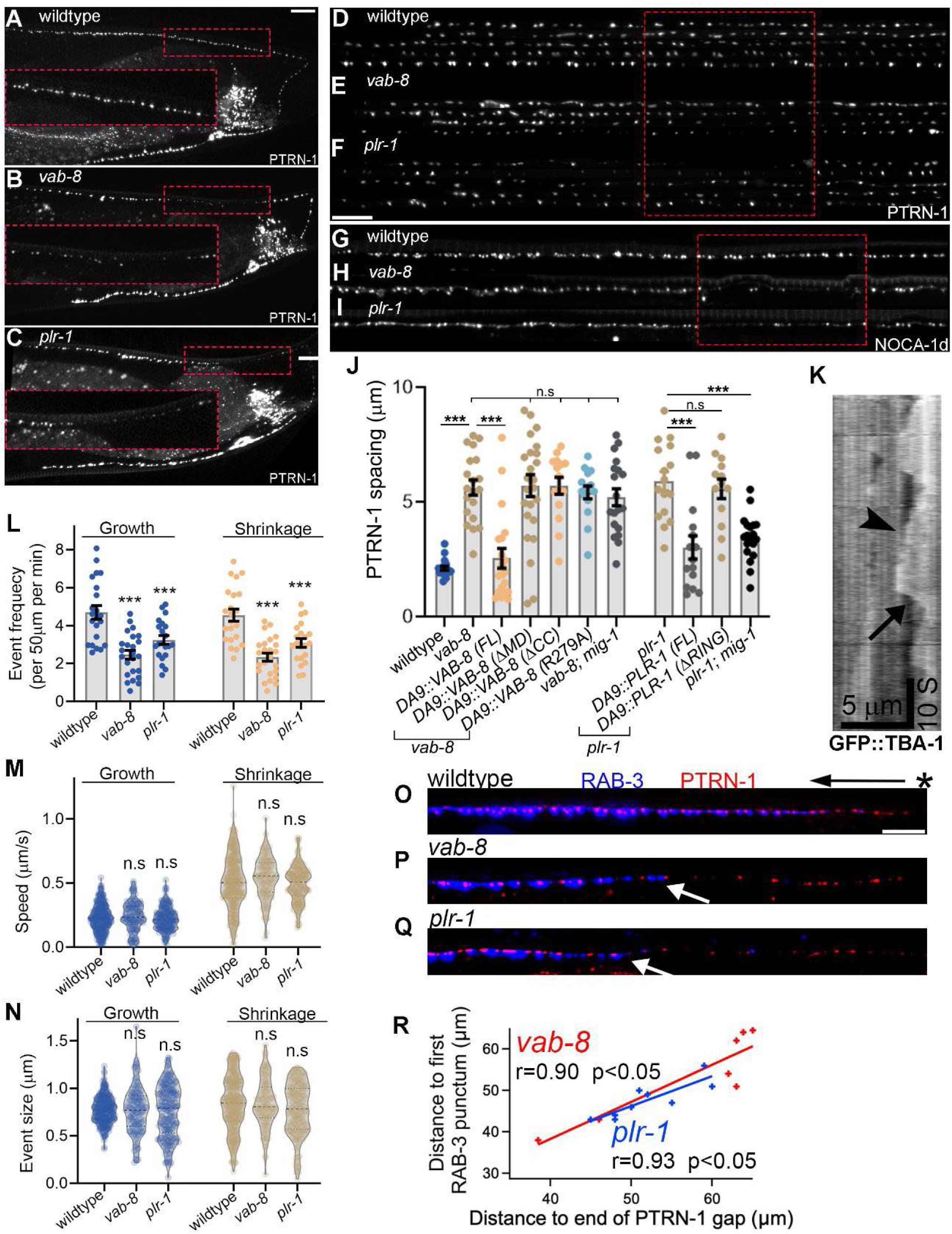
VAB-8 is required for stabilizing minus-end proteins on synaptic MTs. (A-C) Distribution of the MT minus-end marker PTRN-1::YFP in wildtype (A), *vab-8* (B), and *plr-1* mutants (C). Insets show the loss of PTRN-1 from the proximal synaptic region in the mutants. Scale bar 5 μm in all panels. (D-F) Straightened axons from several animals per genotype highlight the stereotypic location where PTRN-1 is lost in the mutants. (G-I) NOCA-1d::GFP is specifically reduced in the proximal synaptic domain of *vab-8* (H) and *plr-1* mutants (I) compared to wildtype (G). (J) Quantification of PTRN-1::YFP puncta spacing in the indicated genotypes. N = 15-25 per genotype. ***p<0.001 (Mann-Whitney U test). (K-N) MT dynamics in the DA9 axon were quantified on kymographs from movies of the MT marker GFP::TBA-1/⍰-tubulin (K). Arrowhead points to polymer growth and arrow points to shrinkage. Overall polymer dynamicity (number of growth and shrinkage events) is reduced in both *vab-8* and *plr-1* mutants (L). The speed of MT growth and shrinkage (M) or the size of individual growth/shrinkage events (N) in the mutants is not significantly different from wildtype. n = 232-539 events from 21-24 animals per genotype. ***p<0.001 (ANOVA with Sidak correction). (O-Q) Co-expression of PTRN-1::YFP and tdTomato::RAB-3 shows at least one MT minus-end in each synapse in wildtype (O). In *vab-8* (P) and *plr-1* (Q) mutants, the severity of RAB-3 phenotypes correlates with the severity of defects in PTRN-1 distribution. (R) Correlation between PTRN-1::YFP gaps and distance to first synapse/tdTomato::RAB-3 in *vab-8* (red) and *plr-1* (blue) mutants. n = 12 (*vab-8*) and 20 (*plr-1*). r Pearson value and statistical significance (Student t test) are shown.

The reduction in PTRN-1::YFP could indicate either a loss of MT polymers in the affected region or a local change in the abundance of minus-end associated proteins. We observed that MTs, labeled with a previously validated GFP::TBA-1/α-tubulin transgene (Yogev et al., 2016), were present in the region where PTRN-1 was reduced (Figure S6), suggesting a specific effect on minus-end proteins. To ask whether dim PTRN-1 puncta in the affected region in *vab-8* mutants still localize to minus-ends, we tested whether the averaged GFP::TBA-1 intensity around them rose to a peak, as would be expected if they are at polymer ends. In wildtype animals, RFP::PTRN-1 puncta were associated with a rise in GFP::TBA-1 intensity, whereas locations midway between RFP::PTRN-1 puncta were not (Figure S7B). A similar pattern was observed in the affected region of both *plr-1* and *vab-8* mutants (Figure S7C-F). These results suggest that VAB-8 is required for the normal accumulation of PTRN-1 on proximal synaptic MTs, but not for the anchoring of MT minus-ends at this location.

To determine the effect of VAB-8 loss on MT organization and dynamics, we imaged animals harboring RFP::PTRN-1 and GFP::TBA-1. *vab-8* and *plr-1* showed fewer MT growth and shrinkage events compared to wildtype, but speed and length of individual events was not affected (Figure 5K-N). Using a method that we had previously developed and validated (Yogev et al., 2016, 2017), we measured average MT length and polymer numbers. Consistent with the observation that PTRN-1 is reduced but not lost in the mutants, we found little effect of *vab-8* or *plr-1* on steady-state MT organization (Figure S7G-L). Although reduced MT dynamicity in the mutants could reflect a reduction in polymer numbers, the lack of effect on GFP::TBA-1 intensity, our MT organization analysis and the fact that PTRN-1::YFP and NOCA-1::GFP intensity are only reduced but not eliminated argue that polymer numbers are not reduced.

### VAB-8 function at MT minus ends is related to its synaptic function

To ask if VAB-8 function at MT minus-ends is related to its synaptic function, we first tested whether both functions require the same protein domains. Cell-specific rescue experiments showed that both VAB-8 and PLR-1 act to maintain the normal distribution of PTRN-1 through the same domains that are required for synapse formation. (Figure 5J). Furthermore, *vab-8* and *plr-1* mutants showed the same genetic interactions with each other and with *mig-1* that were observed for the synaptic phenotype (Figure 5J). Finally, we tested whether the MT and synaptic phenotypes correlate in their severity in individual animals. Co-labeling of PTRN-1 and RAB-3 in *vab-8* and *plr-1* mutants revealed a strong correlation between the extent of synapse-loss and minus end defects (Figure 5O-R). These results suggest that VAB-8 function at MT minus ends is related to its synaptic function.

One possibility is that loss of synapses is the primary defect, which then impairs the recruitment of PTRN-1 to minus-ends. To test this, we examined mutations in kinesin-3/KIF1A *unc-104 (e1265)*, in which DA9 synapses are largely eliminated (Hall and Hedgecock, 1991; Ou et al., 2010). *unc-104(e1265)* mutants did not show a significant reduction in the recruitment of PTRN-1::YFP to discrete puncta, arguing that synapse loss is not the reason for the loss of PTRN-1 from minus-ends in *vab-8* mutants (Figure S5). We next asked whether reduced PTRN-1 or reduced MT dynamicity could underlie the synaptic phenotype. However, DA9 synapses were largely unaffected in *ptrn-1* mutants or in a β-tubulin/ *tbb-2 (qt1)* mutant that strongly suppresses MT dynamics in DA9 (Yogev et al., 2017) (Figure S5). We conclude that loss of synapses does not impair PTRN-1 recruitment and that reduced PTRN-1 and MT dynamicity do not cause the synaptic phenotype.

### VAB-8 regulates cargo pausing during axonal transport

We next turned to the regulation of axonal transport as a possible link between VAB-8 function at MT minus ends and synapses. Local regulation of synaptic cargo delivery at the minus ends was a plausible idea given that the frequency of minus ends is higher than that of synapses, such that every synapse in wildtype DA9 is associated with at least one MT minus-end (Figure 5O). We previously showed that SVP transport pauses at MT ends (Yogev et al., 2016), raising the possibility that UNC-104 could be regulated at the plus-end and dynein at the minus-end. Consistently, Prolonged pausing by UNC-104 is associated with premature synapse formation in *C. elegans*, and pausing facilitates synaptic cargo retention in cultured hippocampal neurons (Guedes-Dias et al., 2018; Wu et al., 2013).

To test this model directly, we examined pausing of GFP::RAB-3 vesicles during transport in the region affected by *vab-8* mutants. In *plr-1* and *vab-8* mutants, pauses were significantly shorter compared to wildtype (Figure 6E). These results indicate that VAB-8 functions to promote pausing during transport, potentially to facilitate cargo delivery. In addition, we observed that cargo run length was increased in both mutants (Figure 6D). This is expected since measured run length would increase when pauses are shorter than our temporal resolution of 100 ms. Vesicle speed and the ratio of anterograde to retrograde movement did not show a consistent change in the two mutants compared to wildtype. We did not detect movement of the active zone marker CLA-1 in either wildtype or mutants (Figure S8A). These results identify VAB-8 as a regulator of synaptic cargo pausing during transport.

**Figure 6:**
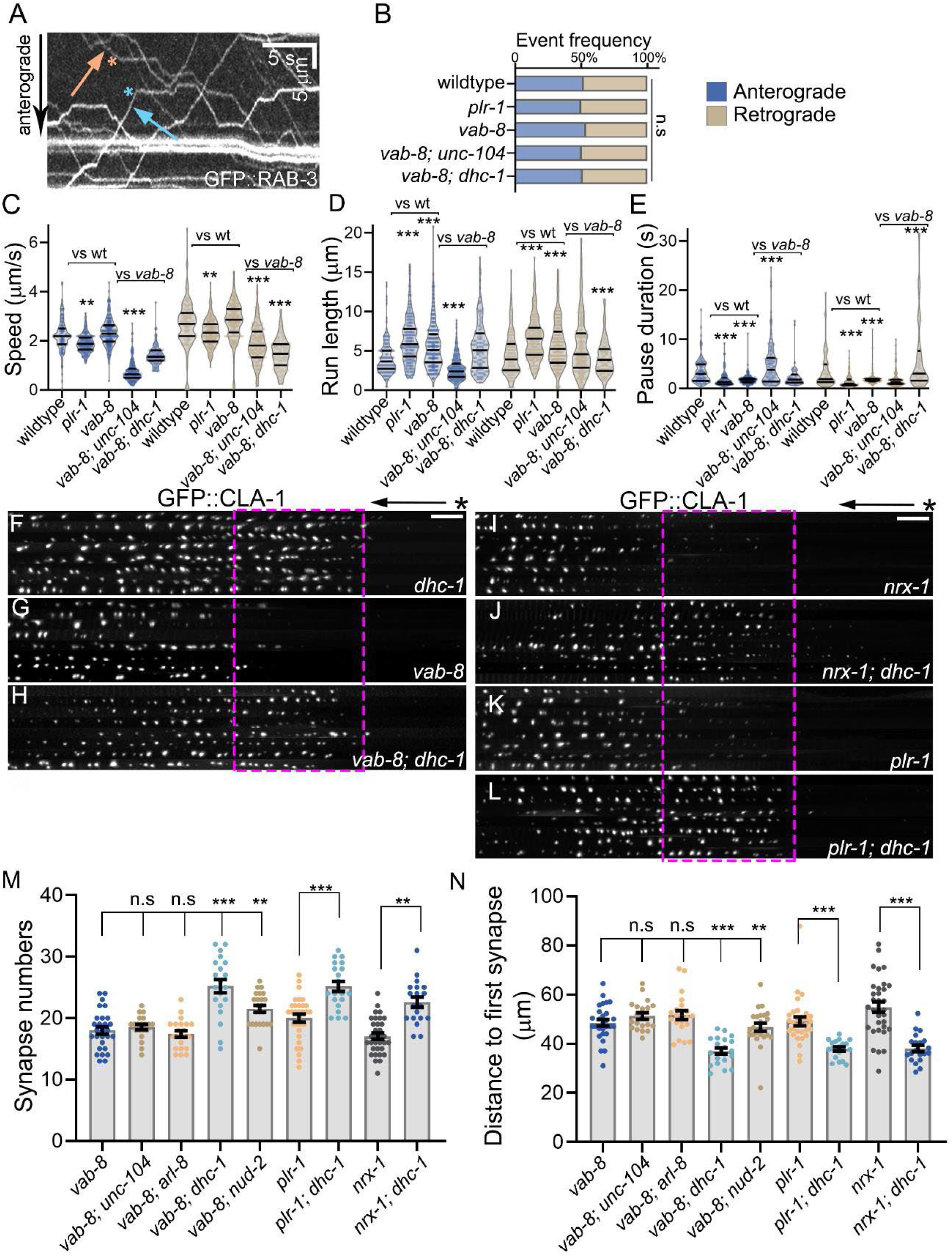
Synapse loss in *vab-8* and *nrx-1* mutants is due to excessive dynein-driven retrograde transport. (A) Example kymograph from a GFP::RAB-3 movie. Arrows and asterisks show runs terminating in pauses. (B-E) Quantification of SVP (GFP::RAB-3) transport in the DA9 axons from indicated genotypes showing overall more processive transport in the *vab-8* and *plr-1* mutants. (B) ratio of anterograde to retrograde events. (C) speed (D) run length and (E) Pause duration. n = 146-520 events from 15-30 animals per genotype. ***p<0.001; **p<0.01; (ANOVA with Sidak correction). (F-L) Reducing dynein function rescues synaptic defects in *vab-8*, *plr-1* and *nrx-1* mutants. Images show 8 axons expressing GFP::CLA-1 aligned to the turn of the commissure. Genotypes: *nrx-1* (H), *nrx-1*; dhc-I (I); *vab-8* (J), *vab-8*; *dhc-I* (K), *plr-1* (L), *plr-1*; dhc-I (M). Scale bar 5 μm. (M-N) Quantification of synapse numbers (M) and distance to first synapse/CLA-1 punctum (N) in the indicated genotypes. n=20-33; ***p<0.001; **p<0.01; (Mann-Whitney U test).

### Synapse loss in *vab-8* and *nrx-1* mutants is due to excessive dynein-driven retrograde transport

To test whether reduced pausing at MT ends is responsible for synaptic defects in *vab-8* mutants, we asked whether compromising the processivity of axonal transport would rescue their synaptic phenotypes. To mildly disrupt transport in the anterograde direction we chose weak alleles of kinesin-3/UNC-104 and its activator ARL-8 (*wy711* and *tm2388*, respectively) (Niwa et al., 2016; Wu et al., 2013) and for retrograde transport we chose a weak allele of dynein heavy chain *dhc-1 (js319)* and its regulator NDEL1 *nud-2 (ok949)* (Koushika et al., 2004; Ou et al., 2010). We confirmed previous results, that these weak mutants do not display overt synaptic defects, with the exception of a mild posterior shift in synapse location in *arl-8 (tm2388)* (not shown) (Ou et al., 2010; Wu et al., 2013).

Double mutants between *vab-8* and *unc-104 (wy711)* or *arl-8 (tm2388)* were similar to *vab-8* single mutants (Figure 6M, N), suggesting that elevated anterograde transport is not responsible for the Vab-8 phenotype. In striking contrast, reducing retrograde transport in *vab-8* mutants, either with *nud-2 (ok949)* or more drastically with *dhc-1 (js319)*, significantly rescued synapse numbers and positioning (Figure 6F, H, M, N). These results suggest that VAB-8 regulates synapse formation and position by antagonizing dynein-driven transport. To test whether the changes we observe in steady-state synapse organization mirror the altered dynamics of axonal transport, we examined vesicle movement in *vab-8 (shy8); unc-104 (wy711)* and *vab-8 (shy8); dhc-1 (js319)* double mutants. Both *unc-104 (wy711)* and *dhc-1 (js319)* have previously been shown to impair transport dynamics in the anterograde and retrograde direction, respectively (Maeder et al., 2014b). In agreement, we found that *unc-104 (w711)* reduced vesicle speed and run length while increasing pause duration in the anterograde direction in *vab-8* mutants. Conversely, *dhc-1 (js319)* reduced retrograde speed and run length while increasing retrograde pauses in *vab-8* mutants. These results support the idea that synapse organization defects in *vab-8* mutants reflect increased retrograde transport.

We next asked whether excessive retrograde transport also underlies synapse loss in *plr-1* and *nrx-1* mutants. Previous live-imaging of cargo transport in *nrx-1* mutants indeed revealed an increase in processivity that is highly reminiscent of our observations in *vab-8* and *plr-1* mutants (Kurshan et al., 2018). *dhc-1(js319)* robustly suppressed the synaptic phenotypes of *plr-1* (Figure 6K-N). *dhc-1(js319)* also strongly suppressed the synapse-positioning defects of *nrx-1* mutants and led to an increase in synapse numbers (Figure 6I, J, M, N). Taken together, our results identify the regulation of retrograde transport at MT minus-ends by VAB-8 as a mechanism through which Neurexin and Frizzled signaling sculpt synaptic connectivity.

Finally, we asked why proximal synapses are specifically affected in *vab-8* mutants. One possibility is that these synapses are more reliant on cargo supply in the retrograde direction. To test this, we photobleached individual proximal and distal boutons in wildtype animals and measured cargo retention events, which we defined as pauses longer than 30 seconds (Figure S8B-D). The rate of retentions was relatively low, consistent with previous studies (Guedes-Dias et al., 2018). Importantly, the number of anterograde and retrograde retention events were similar in each location and both proximal and distal synapses showed equivalent ratios of anterograde to retrograde retentions (Figure S8D). While we cannot completely exclude a differential reliance on anterograde or retrograde transport for cargo delivery, our results argue against this model. Another possibility is that proximal synapse loss in *vab-8* mutants reflects the activity of local anti-synaptogenic signals, consistent with the fact that several extracellular cues restrict the location of DA9 synapses (Klassen and Shen, 2007; Mizumoto and Shen, 2013b, 2013a; Poon et al., 2008). We hypothesized that such signals may also affect nearby neurons, particularly DB7, a motor neuron that synapses in the same region as DA9 but projects its axon posteriorly, such that the location of affected proximal synapses in DA9 corresponds to the distal synapses in DB7 (Figure S8E, F). In DB7, *vab-8* mutants showed specific loss of distal synapses (Figure S8G-I), consistent with our hypothesis. Furthermore, synapse loss in DB7 was suppressed in *vab-8; dhc-1* double mutants, suggesting a similar underlying cell-biological mechanism to DA9 (Figure S8G-I). These results suggest that the local effect of *vab-8* mutants does not reflect a unique property of proximal synapses but likely the effect of signaling pathways that locally control synapse formation.

## Discussion

How the synaptogenic activity of cell-surface molecules promotes delivery of synaptic cargo in transport is poorly understood. We found a role for VAB-8/KIF26 in linking Neurexin and Frizzled signaling to the delivery of retrogradely transported synaptic cargo. We find that the levels of VAB-8 on MT minus-ends in a subset of presynaptic sites are regulated by MIG-1/Fz and Neurexin signaling. In *vab-8* mutants, or when VAB-8 levels are locally reduced in *plr-1* and *nrx-1* mutants, those specific synapses are lost. Conversely, VAB-8 overexpression rescues synapse-loss in *plr-1* and *nrx-1* mutants, suggesting that it is an important mediator of PLR-1’s and Neurexin’s presynaptic function. In *vab-8* mutants, *nrx-1* mutants (Kurshan et al., 2018) and *plr-1* mutants, retrograde axonal transport becomes overly processive. Importantly, synapse loss in these mutants can be rescued by compromising retrograde transport. Together, these results support a model in which the pro- and anti-synaptogenic activities of Neurexin and Fz converge on VAB-8/KIF26 to regulate the delivery of synaptic cargo undergoing retrograde transport from MT minus-ends. This model is further supported by the localization of both paused dynein (Yogev et al., 2016) and VAB-8/KIF26 to MT minus ends.

### Synaptogenic activity promotes the local delivery of cargo undergoing transport

Synapse formation entails the recruitment of actin, scaffolding and active zone molecules, and vesicle precursors (Rizalar et al., 2021; Südhof, 2018). Binding partners for Neurexin and Frizzled intracellular domains – including cytoskeletal proteins – have been identified, but how these signaling molecules regulate presynapse formation is not fully understood. (Biederer and Südhof, 2000; Biederer and Sudhof, 2001; Dean et al., 2003; Hata et al., 1996; Mukherjee et al., 2008; Muhammad et al., 2015; Owald et al., 2012; Lüchtenborg et al., 2014; Miech et al., 2008; Sugie et al., 2015.). Our results identify the regulation of cargo delivery during retrograde transport as a facet of Neurexin’s and Frizzled’s synaptogenic activity: synapse loss in *nrx-1* and *plr-1* mutants is rescued by VAB-8 overexpression or by reducing retrograde transport. How Neurexin and Frizzled signal to VAB-8 remains to be elucidated. Proteomic screens identified a physical interaction between homologs of VAB-8 and LIN-2 or LIN-7 (Hein et al., 2015; Huttlin et al., 2017, 2021) which can bind Neurexin’s cytoplasmic tail (Südhof, 2017), suggesting a plausible mechanism. We have seen mild synaptic phenotypes in *lin-7* mutants, which were not identical to *vab-8* mutants, suggesting that additional mechanisms may be involved (not shown). Previous work identified proteasomal degradation as a mechanism to regulate Kif26B levels during kidney development and in response to Wnt/ROR signaling (Susman et al., 2017; Terabayashi et al., 2012). Proteasomal degradation may also mediate the local loss of VAB-8::GFP from the synaptic region in *plr-1* mutants, but additional work will be needed to identify the relevant degradation pathway.

### Transport pausing at MT minus-ends facilitates cargo capture by synapses

How synaptic material is distributed among *en passant* boutons is poorly understood. Live imaging of DCVs revealed delivery during both anterograde and retrograde transport (Bharat et al., 2017; Cavolo et al., 2016; Wong et al., 2012). Recent work elucidated a mechanism for cargo delivery during anterograde transport: dissociation of kinesin-3/KIF1A from GTP-rich dynamic MTs at synapses (Guedes-Dias et al., 2018). Our single bouton FRAP supports the model of bidirectional delivery of cargo and extends it to include SVPs in *C. elegans*. Furthermore, our results identify MT minus-ends as sites for cargo delivery by dynein during retrograde transport and illustrate how synaptogenic signals converging on VAB-8 can regulate this process. This mode of cargo delivery is not dedicated to proximal or distal boutons, since in DA9 it affects proximal boutons whereas in DB7 the distal boutons are affected (Figure S8).

Recent work provides insight into synaptic capture mechanisms for transport cargo. They include interactions with scaffolding proteins and phosphorylation of SVP proteins (Bharat et al., 2017; Stucchi et al., 2018). These capture mechanisms are likely inefficient (Guedes-Dias et al., 2018), which may be important for distributing cargo rather than leaving it at the first synapse encountered during transport. Hence, we propose that the time a vesicle stays paused on a MT end in the synaptic region is important to allow its capture. The reduced pause duration in *vab-8* mutants, as well as the correlation between longer pauses in *vab-8; dhc-1* double mutants and the rescue of the synaptic phenotype, are consistent with this interpretation.

### VAB-8/KIF26 function at MT minus ends

How does VAB-8 regulate the accumulation of minus-end proteins at synapses? In vitro VAB-8 does not directly bind minus-ends, suggesting that it is recruited to these locations in vivo, at least partially by NOCA-1. Since VAB-8 is found on most, if not all, minus ends in axons and dendrites, its local functions likely reflect the activity of signals operating in the synaptic region. MIG-1/Fz contributes to this local effect, but other signals which remain to be identified likely exist. This is suggested by the fact that in *plr-1; mig-1* double mutants, in which the distribution of synaptic cargo is rescued, VAB-8::GFP is not completely restored to the synaptic region. Furthermore, we observed that in *nrx-1* mutants, although VAB-8 is reduced in synapses, the distribution of PTRN-1::YFP and NOCA-1::GFP was unaffected (not shown). This suggests that a mere reduction in VAB-8 is not sufficient to affect PTRN-1 and NOCA-1, and that a second signal needs to be activated. Complete loss of VAB-8 (as opposed the mere reduction that is observed in *nrx-1* mutants) may trigger activation of the signal that locally disrupts PTRN-1 and NOCA-1. This interpretation is consistent with previously described roles for VAB-8 and KIF26 in regulating signaling pathways such as Netrin, Slit, FAK and GDNF (Wang et al., 2018; Zhou et al., 2009).

## RESOURCE AVAILABILITY

### Lead contact

Further information and requests for reagents should be directed to Shaul Yogev.

### Materials Availability

All transgenic *C. elegans* strains generated for this study are available from the lead contact upon request.

### Data and Code Availability

Code and raw data used for this study are available from the lead contact upon request.

## EXPERIMENTAL MODEL AND SUBJECT DETAILS

### Strains and maintenance

All *C. elegans* strains were cultured on nematode growth medium plates seeded with *E. coli* OP50 as previously described (Brenner, 1974). Animals were grown at 20°C prior to experiments/injections and at room temperature afterward. The N2 Bristol strain was as wildtype for outcrosses.

## METHOD DETAILS

### Genetic screen

We performed a visual forward F2 genetic screen of ∼ 2,000 haploid genomes. Ethyl methanesulfonate (EMS) was used to induced random germline mutations in animals carrying the *wyIs802 [Pitr-1::PTRN-1::YFP]* marker. Mutagenized worms were scored for changes in YFP::PTRN-1a in DA9. Homozygous mutants were rescued and outcrossed 5 times with N2 males prior to phenotypic analysis and whole-genome sequencing. Genomic DNA was purified with Phenol/CHCl_3_ (Invitrogen-Thermo Fisher) and precipitated with ethanol/ NaCH_3_COO (Sigma-Aldrich). DNA was resuspended and solubilized in DNAse-free water (Sigma-Aldrich) and sequenced on Illumina Hi-Seq machines at the Yale Center for Genome Analysis. Sequencing data was analyzed by using tools on Galaxy Project. Briefly, reads were aligned against *C. elegans* N2 genome as a reference. Variants were called and filtered to isolate strain-specific single-nucleotide and indel mutations. The causality of mutations in *vab-8* and *plr-1* for the phenotypes was confirmed by rescue experiments, testing additional alleles and generation of additional mutants using CRISPR.

### Cloning and constructs

All plasmids were generated by Gibson Assembly and confirmed by Sanger sequencing. pSM backbone was used for worm expression plasmids and pMT_mNeonGreen for expression in S2 cells. cDNA was prepared from *C.elegans* lysates according to standard procedures. Sequences and plasmids are freely available upon request.

### CRISPR/Cas9 genome editing

We used CRISPR to generate *shy72* and *shy73* mutations in *vab-8* near the *shy8* mutation in order to confirm the causality of mutations in that region to the *Vab-8* phenotype and in order to produce double mutants *vab-8, nrx-1* since the two genes are too close to each other for generating recombinants. We followed the protocol of (Dokshin et al., 2018). Cas9 protein was purchased from IDT. sgRNA was synthesized using EnGen kit from NEB using the following primer: ttctaatacgactcactatagTCAATCCCTCCCATGCTCCGgttttagagctaga. The following single strand repair template was used to induce STOP mutations: AACACACAACTGCCACGACGGTTGTATTCATTCAATCCCTCCCATGCTATAAGGATCCCTAATTAACCGTAGACACACGCCATTCCTCTCTGCATCACTGAAACTGTACGACGA. Sanger sequencing was used to confirm the repair.

### Fluorescence microscopy

Animals were synchronized as L4 larvae and grown for an additional 20-24 hrs at 20 degrees to reach adulthood. We immobilized worms on agar pads with 5-10 mM Levamisole in M9 buffer for standard fluorescence imaging. Images were acquired through an HC PL APO 63x/1.40NA OIL CS2 objective or a PL APO 40x/1.30NA CS2 objective built on a Laser Safe DMi8 inverted microscope (Leica) equipped with a VT-iSIM system (BioVision). Images were captured with an ORCA-Flash4.0 camera (Hamamatsu) and controlled by MetaMorph Advanced Confocal Acquisition Software Package. For any given set of data, the same settings (laser power, exposure time, and gain) were used.

For time-lapse imaging, animals were incubated in levamisole (0.5 mM) in M9 at room temperature for 5 min prior to mounting on 10% agarose slide in M9 without paralytics, ensuring a low concentration of levamisole that was determined not to produce artifacts. Movies were collected on an Olympus BX61 microscope equipped with a Hamamatsu ORCA-Flash4.0 LT camera at 3 or 10 Hz (microtubules and synaptic vesicles respectively) for 3 min. Image acquisition was monitored by Volocity 2.0 (Quorum technologies). For any given set of data, the same settings (laser power, exposure time, and gain) were used. FRAP of GFP::RAB-3 and VAB-8::GFP was achieved by a 50 milliseconds pulse of 488-laser with a Perkin-Elmer Photokinesis unit.

### DA9 behavior assay

DA9 optogenetic stimulation was performed as previously described (Ding and Hammarlund, 2018). Worms expressing Chrimson in DA9 were cultured on NGM plates seeded with OP50 containing 100 μM all-trans-retinal (ATR). L4s progenies were transferred to a fresh plate without bacteria for the behavior assay. The plate was placed on a Leica M165FC stereo scope, and a Basler acA2440 camera controlled by the WormLab software was used to record movies. The movies were 35 seconds long (10 seconds of prestimulation, 5 seconds of light stimulation and 20 seconds of poststimulation). Bright field illumination was kept at low intensity throughout the recording to avoid any non-specific activation of Chrimson. Continuous green light stimulation was delivered by a CoolLED pE-300^white^ light source. The tail bending behavior was later analyzed in the Wormlab software. In brief, an animal was detected by thresholding, and then the body midline was extracted and divided evenly into 13 segments with 14 points. The angle between the last two segments was calculated as the tail angle for each frame. For frames where the worm contour cannot be detected accurately by the software, we manually drew the body midline.

### Cell culture, transfection, and lysate preparation

*C. elegans* proteins (VAB-8, NOCA-1, PTRN-1) were expressed in *Drosophila* S2 cells whereas rat KIF5C(1-560-mNG) was expressed in COS-7 cells. S2 were cultured in Schneider’s *Drosophila* medium (Gibco) supplemented with 10% (vol/vol) FBS (HyClone) at 26°C and transfected using Lipofectamine LTX with PLUS reagent (Invitrogen). Protein expression was induced by adding 1mM CuSO_4_ to the medium 4-5 h after transfection and the Halo tag was fluorescently labeled by the inclusion of 50 nM JF552/JF646 Halo ligand (Janelia Farms) in the growth medium. COS-7 (African green monkey kidney fibroblasts, American Type Culture Collection, RRID:CVCL_0224) were grown at 37°C with 5% (vol/vol) CO_2_ in Dulbecco’s Modified Eagle Medium (Gibco) supplemented with 10% (vol/vol) Fetal Clone III (HyClone) and 2 mM GlutaMAX (L-alanyl-L-glutamine dipeptide in 0.85% NaCl, Gibco). COS-7 cells were transfected using TransIT-LT1 transfection reagent (Mirus). Cells are checked annually for mycoplasma contamination.

To prepare cell lysates for single-molecule assays, cells were harvested 24 h (COS-7) or 48 h (S2) post-transfection and pelleted by low-speed centrifugation at 4°C. The cell pellet was washed with PBS buffer and resuspended in ice-cold lysis buffer (25 mM HEPES/KOH, 115 mM potassium acetate, 5 mM sodium acetate, 5 mM MgCl_2_, 0.5 mM EGTA, 1% Triton X-100, pH 7.4) freshly supplemented with 1 mM ATP, 1 mM PMSF, 1 mM DTT and protease inhibitors (Sigma-Aldrich). The cell lysate was clarified by centrifugation at full-speed at 4°C and aliquots of the supernatant were snap frozen in liquid nitrogen and stored at -80°C.

The amount of VAB8(1-795)::mNG and VAB8(1-795,R279A)::mNG in the S2 cell lysates was quantified by a dot-blot in which a dilution series of S2 lysates was spotted onto a nitrocellulose membrane, air-dried for 1 h, and immunoblotted with a monoclonal primary antibody against mNeonGreen (Chromotek) at room temperature for 80 min and secondary antibody 680nm-anti rabbit (Jackson ImmunoResearch Laboratories Inc.) at room temperature for 1 h. The fluorescent spots were detected by Azure c600 (Azure Biosystems) and their intensity was quantified using Fiji/ImageJ (NIH) to normalize the motor concentrations.

### Immunofluorescence and spinning disk confocal microscopy

Transfected S2 cells were fixed with 3.7% formaldehyde in PBS, permeabilized with 0.2% Triton X-100 in PBS, and then blocked in blocking solution (0.2% fish skin gelatin in PBS). Primary and secondary antibodies were applied in blocking solution at room temperature for 1 h each. Nuclei were stained with DAPI. The glass coverslips were mounted in ProlongGold (Life Technologies). Primary antibodies: mouse anti-β-tubulin (E7, Developmental Studies Hybridoma Bank) and rabbit anti-Halo (G9281, Promega). Secondary antibodies: 594 nm anti-rabbit and 680 nm anti-mouse (Jackson ImmunoResearch Laboratories). Images were collected on an inverted spinning disk confocal microscope (Nikon CSU-X1) equipped with a 100x, 1.49 numerical aperture (NA) oil-immersion objective, four diode lasers (405nm, 488 nm, 561 nm and 647 nm), and an EMCCD detector (iXon Ultra 888, Andor). Z-step series were acquired with a z step per image of 500 nm. A focal plane through the mid-nucleus was chosen for display and to directly compare protein localization.

### Single-molecule motility assays

Microtubules were polymerized from purified tubulin and 10% Hily647-or Hilyte488-labeled tubulin (Cytoskeleton) in BRB80 buffer (80 mM Pipes/KOH pH 6.8, 1 mM MgCl_2_, and 1 mM EGTA) supplemented with 1 mM GTP and 2.5 mM MgCl_2_ at 37°C for 30 min. 20 μM taxol in prewarmed BRB80 buffer was added and incubated at 37°C for additional 30 min to stabilize microtubules. Microtubules were stored in the dark at room temperature. A flow cell (∼10 μl volume) was assembled by attaching a clean #1.5 coverslip (Fisher Scientific) to a glass slide (Fisher Scientific) with two strips of double-sided tape.

Polymerized microtubules were diluted in BRB80 buffer supplemented with 10 μM taxol and then were infused into a flow cell and incubated for 5 min at room temperature for nonspecific adsorption to the coverslip. Subsequently, blocking buffer (1 mg/ml casein and 10 μM taxol in P12 buffer) was infused and incubated for 5 min. Finally, cell lysate diluted in motility mixture [2 mM ATP, 3 mg/ml casein, 10 µM taxol, and oxygen scavenging system (1 mM DTT, 1 mM MgCl2, 10 mM glucose, 0.2 mg/ml glucose oxidase, and 0.08 mg/ml catalase) in P12 buffer] was added and the flow-cell was sealed with molten paraffin wax. Images were acquired by total internal reflection fluorescence (TIRF) microscopy using an inverted microscope Ti-E/B (Nikon) equipped with perfect focus system (Nikon), a 100× 1.49 NA oil immersion TIRF objective (Nikon), three 20-mW diode lasers (488 nm, 561 nm, and 640 nm) and an electron-multiplying charge-coupled device detector (iXon X3DU897; Andor Technology). Images were acquired continuously at 200 ms per frame for 40 s. Image acquisition was controlled using Nikon Elements software and all assays were performed at room temperature. Maximum-intensity projections were generated using Fiji/ImageJ (NIH). To determine whether R279A mutation disrupts the microtubule binding ability of VAB-8, equal amount of VAB-8(1-795)::mNG or VAB-8(1-795,R279A)::mNG protein diluted in motility mixture was added to flow cells. The number of motors on a microtubule was counted and then divided by the length of the microtubule and the recording time to obtain a binding rate with the units of events/μm/min. The binding rate of an adjacent region that lacks microtubules was subtracted as background. Statistical analysis was performed using a two-tailed t-test (****, P<0.0001) and graphs were generated using Prism software (GraphPad). WT, n=35 microtubules and R279A, n=28 microtubules across 3 independent experiments.

## QUANTIFICATION AND STATISTICAL ANALYSIS

### Fluorescence quantification

Raw data were exported for analysis on ImageJ or MATLAB. Kymographs were generated using the ImageJ KymoResliceWide plugin following correction for animal movement using the StackReg plugin. Traces on kymographs were analyzed manually and data was exported to Excel and Igor Pro 8 for further analysis. To quantify the number of synapses, a line-scanning method along the axon was used to obtain the GFP::CLA-1 fluorescent profile. We used the same criterion as (Kurshan et al., 2018) to count a peak in the signal as a synapse (>5% of the maximum) for consistency with previous literature. Note that this criterion usually leads to the scoring of the fainter CLA-1 puncta in *nrx-1* mutants (highlighted Figure 1) as synapses. VAB-8::GFP axonal localization was resolved by drawing a region of interest (ROI) around the axon. Based on the intensity histogram, a default threshold was applied to the ROI to create a binary mask. The mask was firstly treated with the built-in Watershed segmentation algorithm to better define nearby puncta. Lastly, the number of puncta and their intensity were assessed by using the Analyze Particle function provided in ImageJ. As an alternative method for extracting puncta size and intensity (for example Figure 4 and Figure S1) we used the Fiji plugin FindFoci.

PTRN-1 puncta spacing was quantified by measuring the distance between neighboring YFP::PTRN-1a puncta in the axonal region of interest. A custom MATLAB code (available upon request) was used to analyze the distribution of GFP::TBA-1 around RFP::PTRN-1 peaks. Briefly, the code determines the location of PTRN-1 peaks and averages GFP::TBA-1 in a 15-pixel region around each peak. As a control, the midway point between two peaks is used. To average the signal across multiple animals, the “on peak” traces were normalized by “off peak traces” in order to prevent variability in marker expression to impact the analysis. Axonal microtubule organization and minus-end analysis was done using a previously described and validated MATLAB code (Yogev et al., 2016), which is available upon request.

### Statistical analysis

Statistical analysis was performed on Igor Pro 8, Prism 9 (GraphPad), and SPSS (IBM). Data were considered significant at *p* ≤ 0.050 and are plotted as the means with SEM. Each data set consists of at least three independent experiments. Kolmogorov–Smirnov test was used to determine the normality of data, followed by statistical comparisons as noted in figure legends.

---

**Figure S1:**
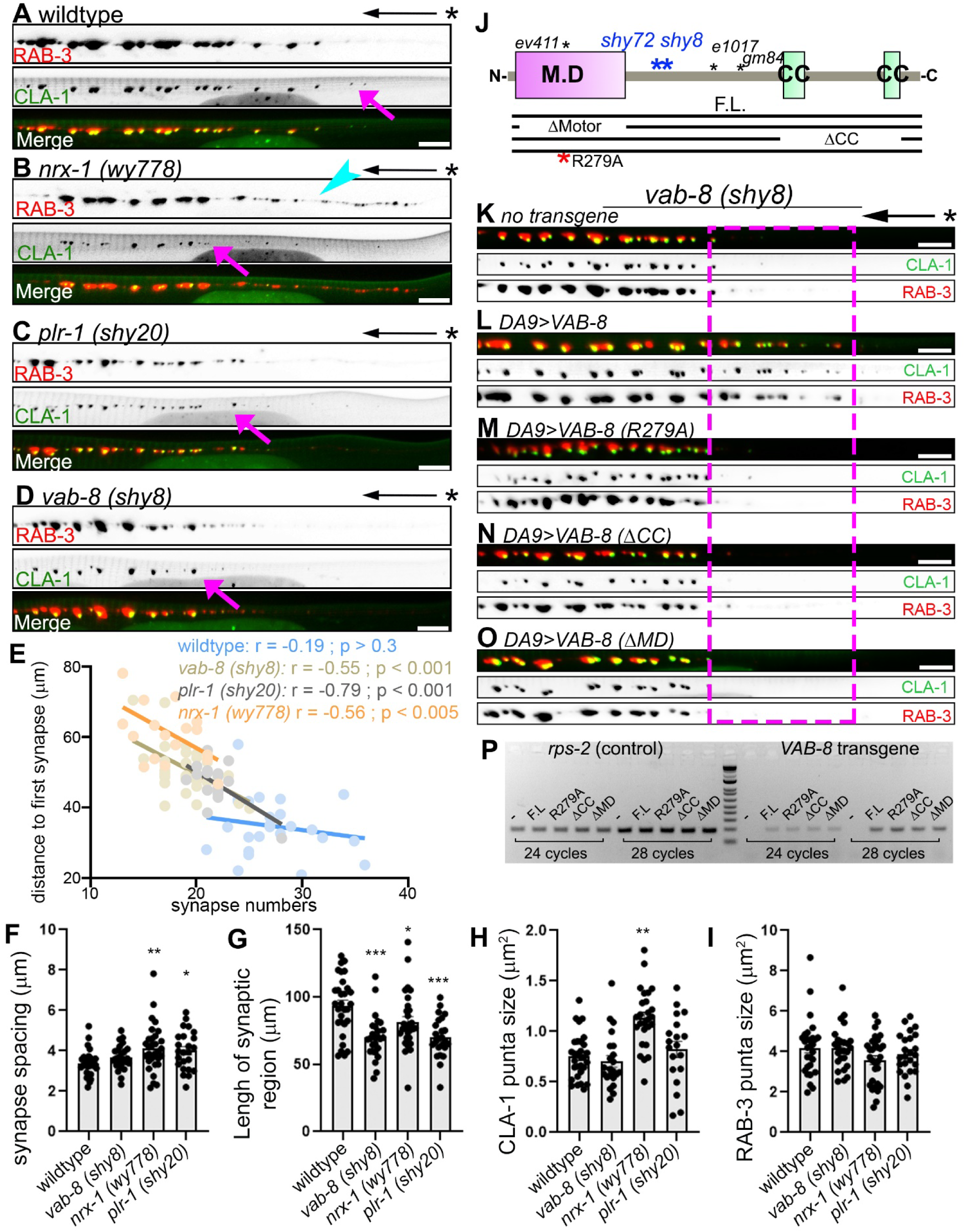
Synaptic phenotypes in *vab-8, plr-1* and *nrx-1* mutants. **(A-D)** *vab-8* and *plr-1* mutants phenocopy the missing proximal synapses of *nrx-1* but do not show vesicle clustering defects. Axons from wildtype (A), *nrx-1* (B), *plr-1* (C) and *vab-8* (D) expressing the active zone marker GFP::CLA-1 and the synaptic vesicle precursor marker tdTomato::RAB-3. * marks the turn of the commissure. Pink arrows point to the first synapse and arrowhead points to RAB-3 that is not associated with CLA in *nrx-1* mutants, indicating vesicle clustering defects. Scale bar = 5 µm. (E) Synapse numbers are inversely correlated to the distance of the first synapse from the commissure in *plr-1 (sh20), vab-8 (shy8)* and *nrx-1 (wy778)* mutants but not in wildtype animals. n=25-34 animals per genotype with Pearson’s r and two-tailed p value shown. **(F-I)** Quantifications of synapse spacing (F), Length of synaptic region (G), CLA-1 puncta size (H), RAB-3 puncta size (I) in the indicated genotypes. n = 24-30 animals per genotype *p<0.05, **p<0.01, ***p<0.001 (ANOVA with Sidak correction). (J) Diagram of VAB-8 domains showing the location of mutations tested and the transgenic rescue constructs tested. M.D: motor domain. CC: coiled coil. **(K-O)** Rescue experiments with VAB-8 deletion constructs. Magenta square highlights region of lost synapses in *vab-8* mutants (K). Full length VAB-8 expressed in DA9 (L) robustly rescues the phenotype. Constructs with a mutation that reduces microtubule binding (L), deletion of the C terminus coiled coil domains (N) or deletion of the motor domain (O) fail to rescue. (P) Semi-quantitative RT-PCR shows comparable expression levels of the rescuing and non-rescuing transgenes. *rps-2* is used as a loading control.

**Figure S2:**
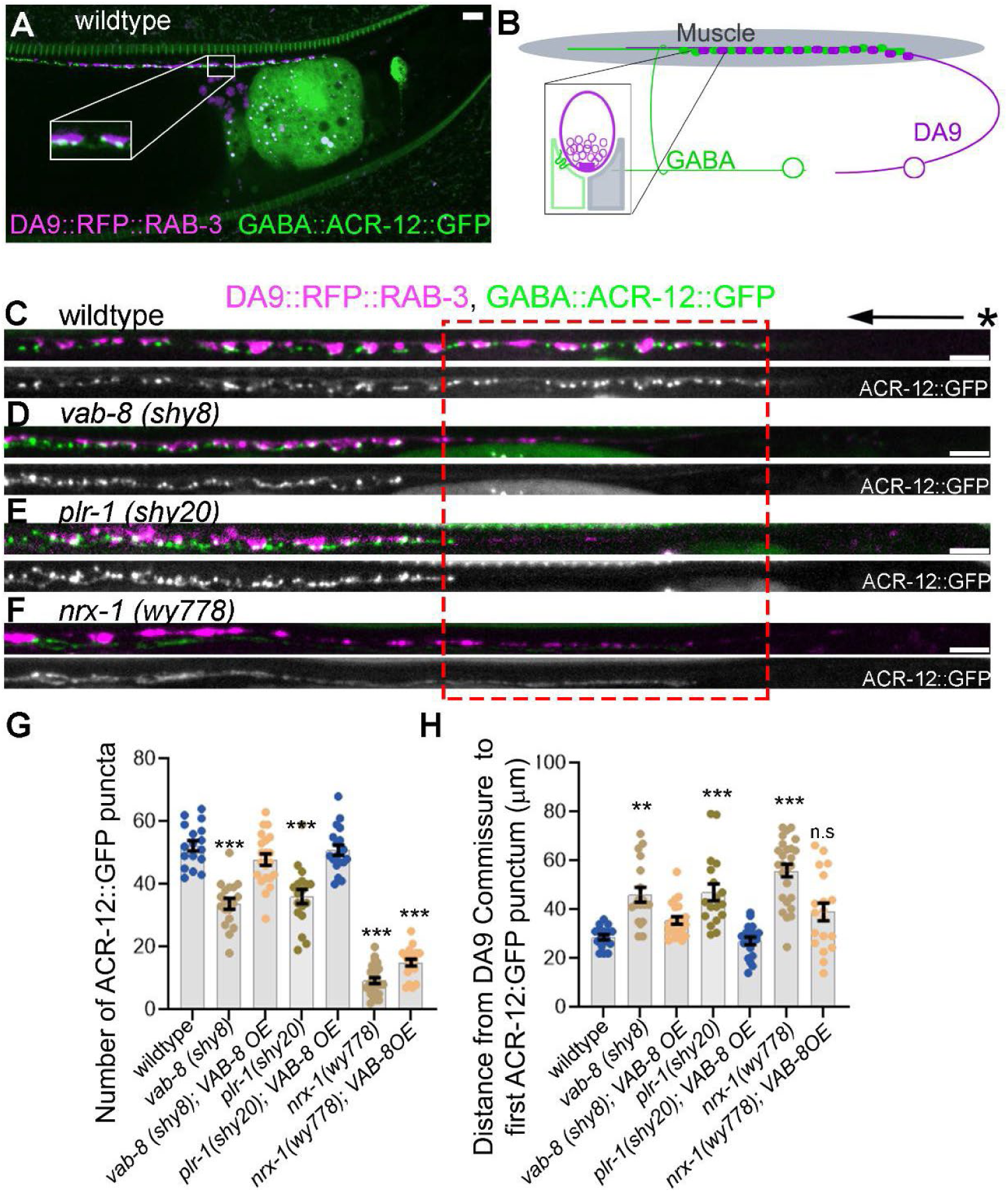
v*a*b*-8 (shy8)* and *plr-1 (shy20)* mutants lack postsynaptic ACR-12. **(A, B)** Confocal micrograph (A) and schematic (B) showing the organization dyadic synapses formed by DA9 onto dorsal muscle and inhibitory GABA neurons. RFP::RAB-3 labels the presynaptic side and ACR-12::GFP labels postsynaptic receptor clusters. Scale bar 5 µm. **(C-F)** Double label confocal images showing the normal apposition of presynaptic RFP::RAB-3 with postsynaptic ACR-12::GFP in wildtype (C), and concomitant loss of proximal pre and post synaptic markers from *vab-8 (shy8)* (D) and *plr-1 (shy20)* (E) mutants. Note that in *nrx-1 (wy778)* mutants ACR-12 fails to cluster, even when presynaptic RAB-3 clusters are present. Scale bar 5 µm. **(G. H)** Quantification of ACR-12::GFP numbers (G) and distance from the DA9 commissure (H) in the indicated genotypes. “VAB-8 OE” denotes presynaptic expression of VAB-8 in Cholinergic neurons using the *unc-17* promoter. n = 18-26, *** p<0.001, ** p<001 (ANOVA with Sidak correction).

**Figure S3:**
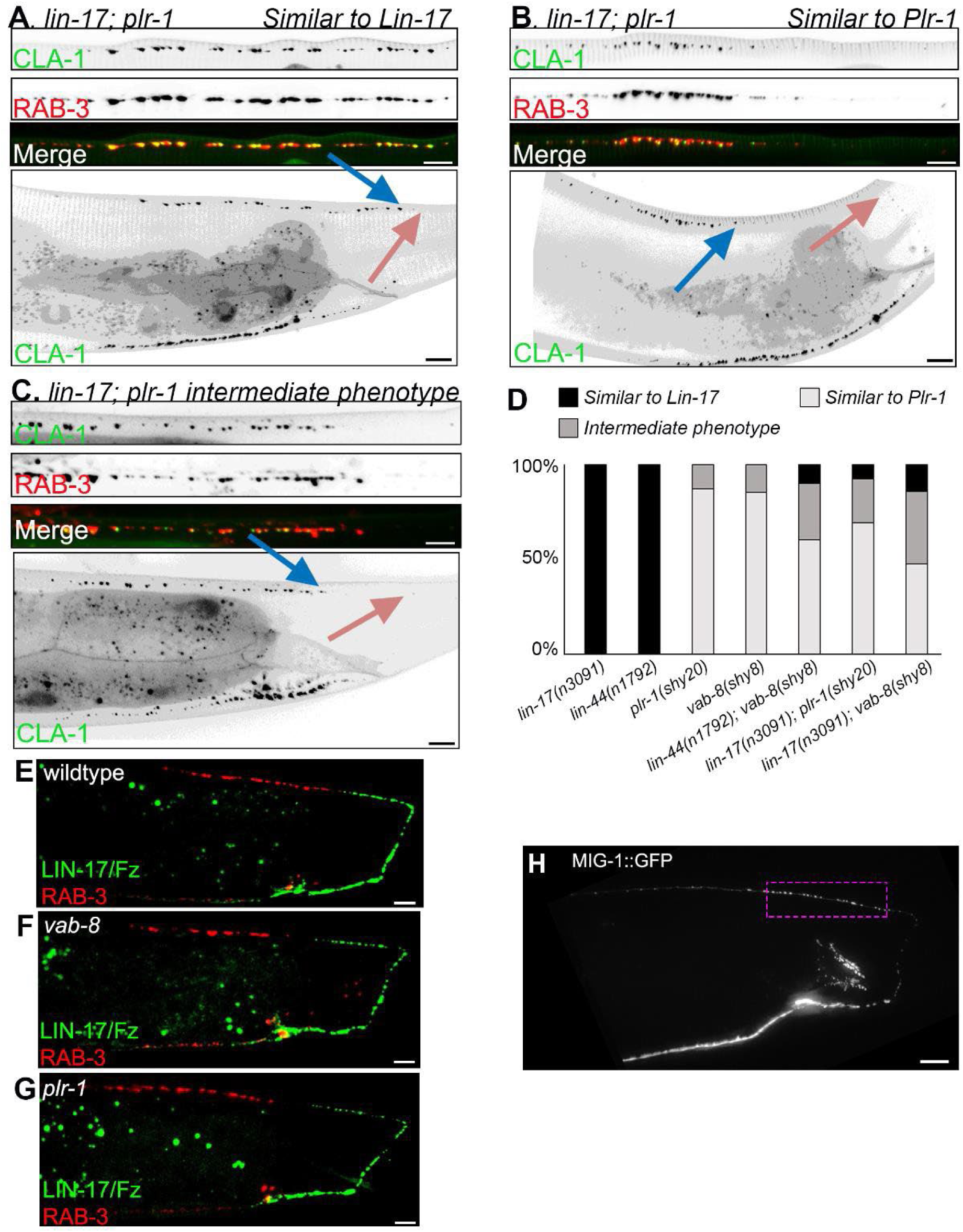
The Frizzled receptor LIN-17 does not act downstream of PLR-1. **(A-C)** Representative images of the different phenotypic classes used for quantifying genetic interactions. All images are from *lin-17; plr-1* double mutants expressing the active zone marker GFP::CLA-1 and the synaptic vesicle precursor marker tdTomato::RAB-3. Pink arrow marks the turn of the commissure and blue arrow marks the first synapse. “*Lin 17 phenotype”* is characterized by synapses starting immediately following the turn of the commissure or in the commissure itself. “*Plr-1 phenotype”* is defined as an asynaptic domain that is larger than in wildtype. “*intermediate phenotype”* is characterized by synapses starting at their normal location or slightly closer to the turn of the commissure. (D) Quantification of the fraction of each phenotypic class in the indicated mutant backgrounds. **(E-G)** Representative images of wildtype (E), *vab-8* (F) and *plr-1* (G) adult worms expressing Frizzled/LIN-17::YFP and mCherry::RAB-3 in DA9. No apparent changes in the localization of Frizzled/LIN-17 is observed in the mutants. Scale bar = 5 µm. (H) Image shows adult worm expressing MIG-1::GFP in DA9. Magenta rectangle indicates the location where synapses are missing in *plr-1* mutants. Scale bar = 5 µm.

**Figure S4:**
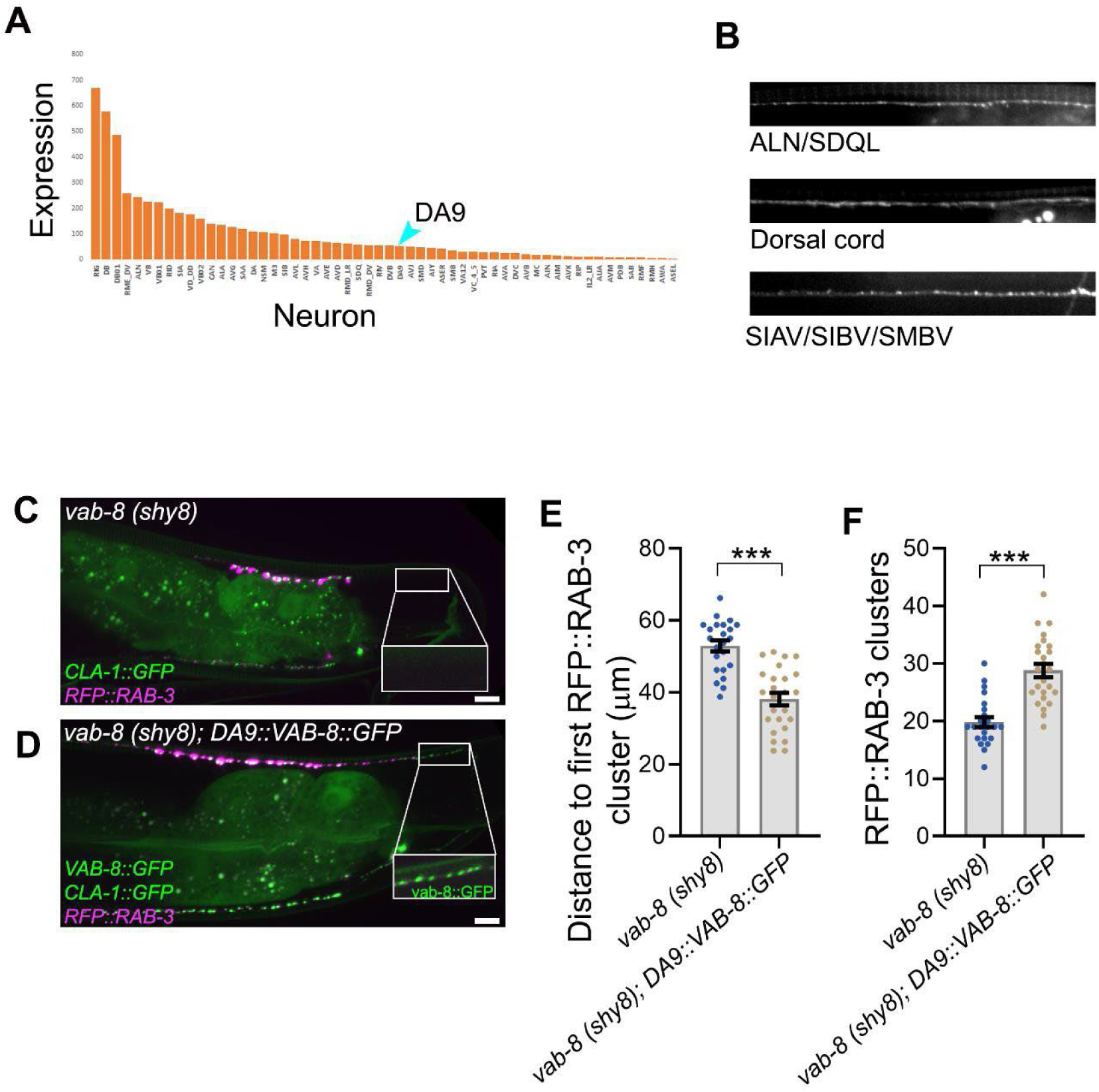
Endogenous VAB-8 expression and VAB-8::GFP rescue. (A) Expression level of VAB-8 in single-cell RNAseq experiments from the CenGen project: https://cengen.shinyapps.io/CengenApp/. DA9 is indicated. (B) select examples of neurons where VAB-8 expression was sufficiently high to be detected with an endogenous 7XspGFP11 tag and pan-neuronal expression of spGFP1-10. Neuron identity was assigned based on the position of the axon (in most cases cell-bodies were too dim) and therefore contains some ambiguity. **(C-F)** GFP-tagged VAB-8 expressed in DA9 with the *Pitr-1* promoter rescues *vab-8* mutants. (C,D) confocal micrograph showing rescue of synapses (RFP::RAB-3, CLA-1::GFP) in *vab-8* mutants expressing *Pitr-1::VAB-8::GFP* (D). Inset shows VAB-8::GFP expression, which can be unambiguously discerned from CLA-1::GFP in asynaptic regions. (E, F) Quantification of synapse number (RFP::RAB-3 clusters, (F)) and distance to first synapse (E) in *vab-8* mutants (n=23) and with rescue transgene (n=25). *** p < 0.001; (Mann-Whitney *U* test).

**Figure S5:**
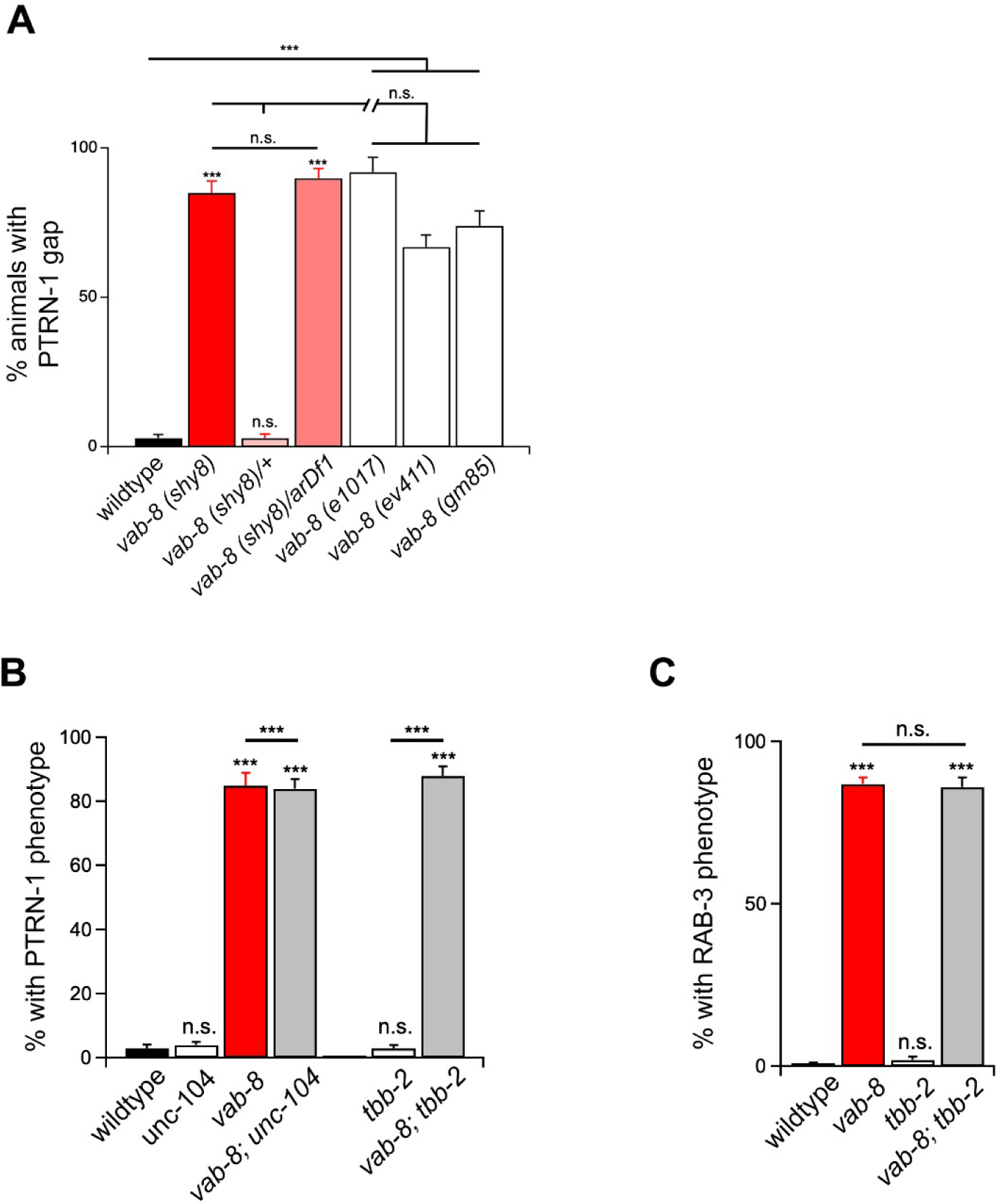
PTRN-1 gaps in *vab-8* mutants are not caused by the loss of synapses and loss of microtubule dynamicity does not lead to synapse elimination. (A) Comparison to previously characterized alleles and a deletion of the *vab-8* region shows that *shy8* is a genetic null allele. The presence of a PTRN-1::YFP “gap” around proximal synapses was scored. (B) The presence of PTRN-1 defects was tested in *unc-104(e1265)* in which DA9 synapses are largely eliminated and in *tbb-2(qt1)* mutants in which microtubule dynamicity is strongly suppressed in DA9. In both cases, no PTRN-1::YFP “gaps” are seen, and double mutants with *vab-8* do not modify the *Vab-8* phenotype, suggesting that loss of synapses and reduced microtubule dynamics do not drive the loss of PTRN-1::YFP in *vab-8* mutants. (C) Suppressing microtubule dynamics in *tbb-2(qt1)* mutants does not lead to loss of proximal synapses and does not modify the loss of proximal synapses in *vab-8* mutants. n = 100-150 per genotype, \*\*\**p*<0.001; (Mann-Whitney *U* test).

**Figure S6:**
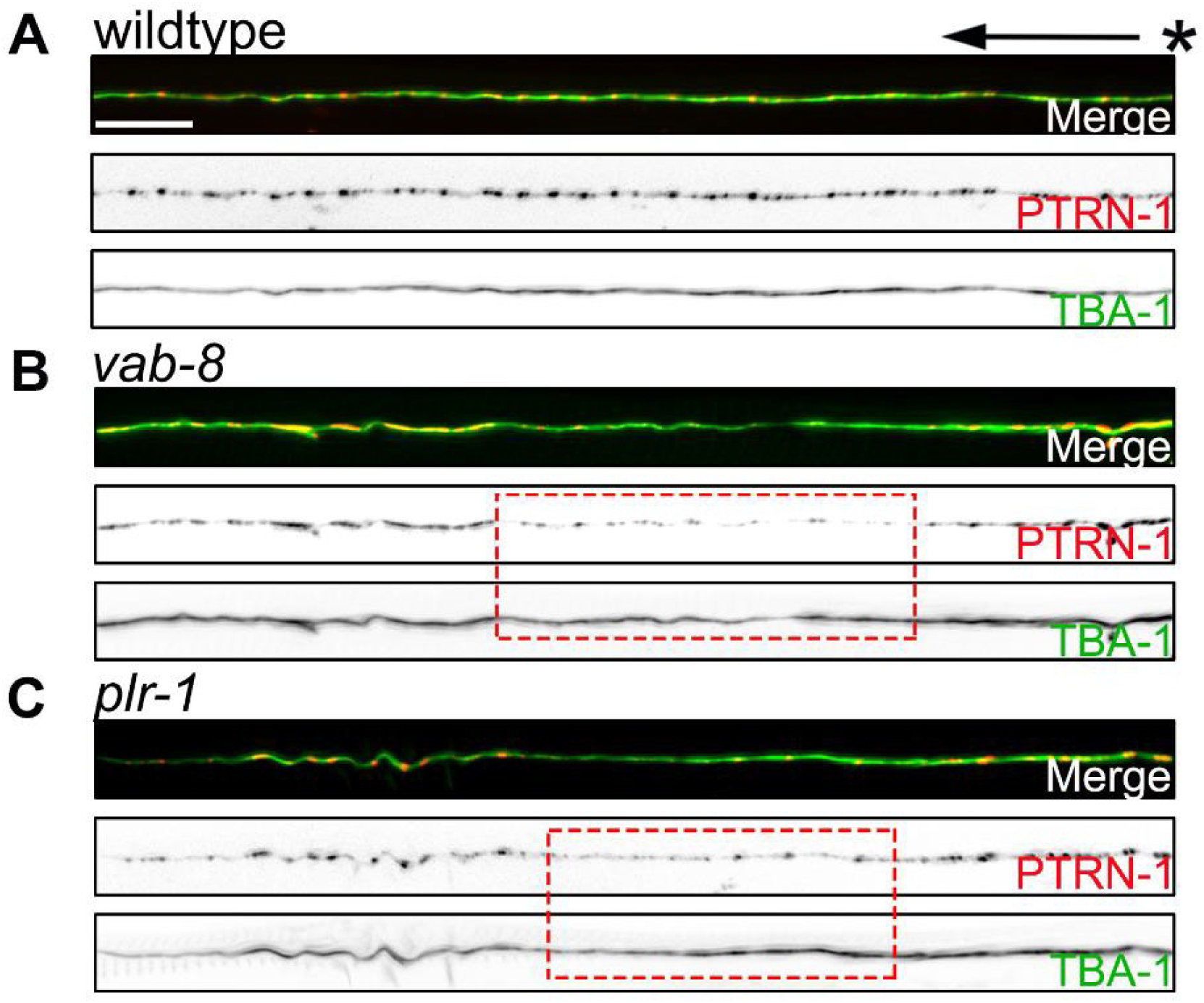
Microtubules are present in the region that shows PTRN-1 gaps in *vab-8* and *plr-1* mutants. **(A-C)** Images of axons from wildtype (A), *vab-8* (B) and *plr-1* (C) mutants expressing RFP::PTRN-1 and GFP::TBA-1/α-tubulin. Loss of PTRN-1 is not associated with loss of microtubules. * marks turn of the commissure. Dashed boxes highlight the affected region. Scale bar = 5 µm.

**Figure S7:**
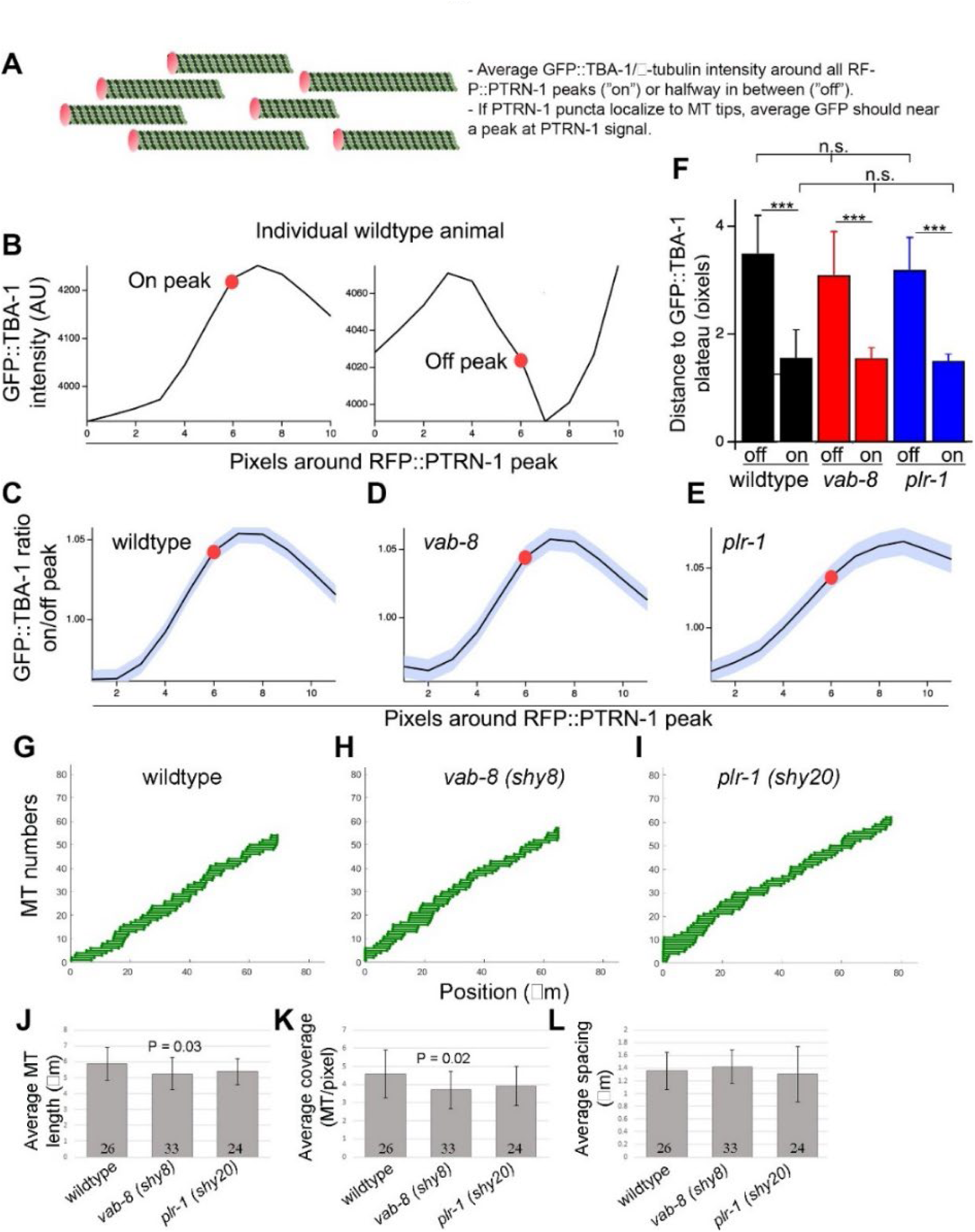
Normal minus-end localization of PTRN-1 and normal microtubule organization in *plr-1* and *vab-8* mutants. (A) Schematic of the concept for determining association of RFP::PTRN-1 with microtubule ends. The intensity of GFP::TBA-1 from an individual microtubule that harbors a given RFP::PTRN-1 punctum cannot be distinguished from that of other microtubules near it and so it is not possible to test where individual PTRN-1 puncta are localized along a given polymer. However, the average GFP::TBA-1 around many RFP::PTRN-1 puncta should increase and near a plateau if the puncta are associated with microtubule ends. (B) Example from a single wildtype animal: average GFP::TBA-1/α-tubulin around all RFP::PTRN-1 puncta from a 45 µm region around proximal synapses shows the expected increase. As a negative control, GFP::TBA-1/α-tubulin shows a decrease around the halfway point between PTRN-1 puncta. **(C-E)** Identical behavior of GFP::TBA-1/α-tubulin around RFP::PTRN-1 peaks in wildtype (C), *vab-8* (D) and *plr-1* (E) confirms that the faint RFP::PTRN-1 puncta in the mutants are associated with microtubule minus-ends. (F) Quantification of the distance between PTRN-1 puncta (“on”) and the halfway between them (“off”) to the nearest peak in GFP::TBA-1/α-tubulin intensity shows no difference between wildtype and mutants. **(G-L)** wildtype animals (G, J), *vab-8* mutants (H,K) and *plr-1* mutants (I,L) show overall similar microtubule organization. Microtubule organization parameters were determined as described in (Yogev et al., 2016). (G,H,I) show models for individual animals. (J) Average polymer length. (K) Average number of polymers per “cross section” of the axon at a given pixel. (L) Spacing between microtubule ends. n = 20-25.

**Figure S8:**
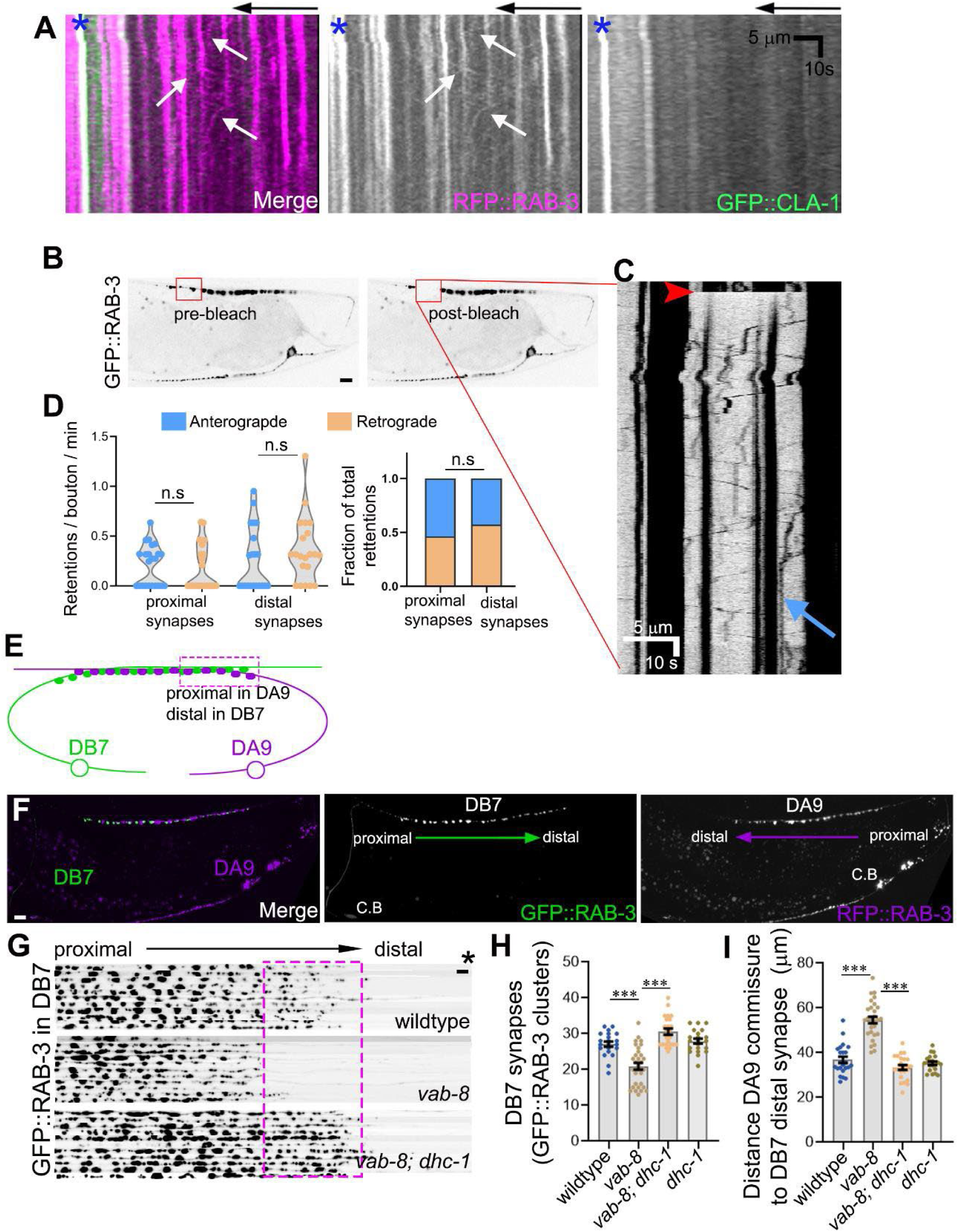
Distal synapses are lost in DB7 neuron in *vab-8* mutants. (A) Example kymograph from a movie showing RFP::RAB-3 and GFP::CLA-1. Arrow indicate movement of RFP::RAB-3 puncta, whereas CLA-1 were stationary in all movies. Arrow indicates the turn of the DA9 commissure and * indicates the proximal-most synapse. (B) Example of FRAP protocol: 1 or 2 synaptic boutons were photobleached either in the distal region (shown) or proximal region (not shown). (C) Example kymograph showing GFP::RAB-3 movement with the distal synaptic region. Red arrowhead indicates photobleaching and blue arrow indicates a retention event. (D) Frequency of GFP::RAB-3 retentions from anterograde/retrograde movements in individual animals (left) and fraction of total retentions from either direction were scored from proximal and distal single bouton FRAP experiments as shown in (C) n=23 and 21 animals for proximal and distal FRAPs, respectively. **(E,F)** Diagram (E) and confocal micrograph (F) showing the spatial organization of DA9 synapses (RFP::RAB-3) and DB7 synapses (GFP::RAB-3). The proximal DA9 synapses are found in the same region as the distal DB7 synapses. Scale bar 5 µm. (G) DB7 synapses (GFP::RAB-3) from 11 animals per indicated genotype were aligned from the turn of the DA9 commissure (indicated with *). *vab-8* mutants show synapse loss, which can be suppressed by *dhc-1* mutants. Note that unlike DA9, where proximal synapses are lost, in DB7 the distal synapses are lost. **(H, I)** Quantification of synapse numbers (H) and distance from DA9 commissure to the distal-most DB7 synapse (I) n=21-29 animals per genotype.

